# Site-directed placement of three-dimensional DNA origami

**DOI:** 10.1101/2022.12.11.519977

**Authors:** Irina Martynenko, Elisabeth Erber, Veronika Ruider, Mihir Dass, Gregor Posnjak, Xin Yin, Philipp Altpeter, Tim Liedl

## Abstract

Assembling hybrid substrates with nanometer-scale precision and molecular addressability enables advances in such distant fields as material research and biosensing. As such, the combination of lithographic methods with 2D DNA origami self-assembly [1–4] has led, among others, to the development of photonic crystal cavity arrays [2] and the exploration of sensing nanoarrays where molecular devices are patterned on the sub-micron scale [5–7]. Here we extend this concept to the third dimension through mounting 3D DNA origami onto nano-patterned substrates followed by silicification [8, 9] to provide mechanical and chemical stability. Our versatile and scalable method relying on self-assembly at ambient temperatures offers the potential to 3D-position any inorganic and organic components that are compatible with DNA architectures [10–13]. This way, complex and 3D-patterend surfaces designed on the molecular level while reaching macroscopic dimensions could supersede energy-intensive manufacturing steps in substrate processing.

## INTRODUCTION

Substrates and surfaces structured on the micron and nanometer scale are ubiquitous in modern life and are used in information technology, bio-sensors, water repellent surfaces or cloth and solar cells. To achieve three-dimensional (3D) architectures in chip technology, for example, multiple lithography steps are executed on top of each other. Replacing top-down lithography in parts or entirely through self-assembly processes could help to reduce production times and energy costs. Structural DNA nanotechnology and in particular DNA origami self-assembly [14, 15] has proven useful in the bottom-up fabrication of well-defined complex designer two-dimensional and three-dimensional nanostructures with single-nanometre feature resolution [16–21]. DNA origami-assisted lithographic methods can successfully transfer spatial information of discrete DNA origami shapes [22–27] or extended 3D periodic DNA lattices [28] into inorganic substrates. Recently, micrometer-scale periodic 3D DNA patterns assembled from DNA bricks [29] were transferred to Si via reactive ion etching, successfully reaching line pitches as small as 16.2 nm, which is already smaller than what is achievable with state-of-the-art quadruple patterning or extreme-ultraviolet lithography [28].

Much of the power of DNA origami lies in its ability to serve as a molecular breadboard for positioning molecules and nanoparticles in space with sub-nm precision [30, 31]. By combining DNA origami self-assembly with lithographic nanopatterning, Gopinath et al. established so-called DNA origami placement (DOP), a technique based on site- and shape-selective deposition of DNA origami objects onto lithographically patterned substrates, creating large-scale arrays of precisely placed DNA structures [1]. DOP overcame some of the drawbacks mentioned above and demonstrated the ultimate power of dictating the nanoscale arrangement of nanocomponents such as metallic nanoparticles [32], organic dyes [2, 4], proteins [33] and peptides [34] over 2D arrays and patterns. Planar triangular or disc-shaped DNA origami were positioned on substrates patterned by e-beam lithography with very high accuracy and orientation control [3, 4]. To further circumvent the complex e-beam steps in this patterning procedure, highly parallel and low-cost methods such as self-assembling nanosphere lithography [35, 36] or nanoimprint lithography [3, 37] of centimeter-sized substrates were recently applied. However, all DOP methods developed so far are limited to planar DNA origami and can fabricate 2D arrays and patterns only.

Herein, we demonstrate site-directed placement of various 3D DNA origami shapes in nanometer-precise patterns over micro-to millimeter scales. We employed two different approaches to achieve the up-right positioning of various DNA origami shapes via connector-mediated binding (hollow tubes) or direct binding (barrels, tetrapods) via self-aligning. Both approaches are compatible with the two nanopatterning techniques that we tested, e-beam lithography and nanosphere lithography. To mechanically and chemically stabilize the arrays, the DNA structures were silicified on their respective substrates resulting in hybrid DNA–silica structures with controllable heights up to 50 nm and a feature size down to ~ 6 nm. Finally, as a proof of concept, we connected the individually placed DNA origami in the xy-plane with further DNA struts in order to create continuous periodic networks.

## RESULTS AND DISCUSSION

The various steps of the fabrication process are illustrated in Figure 1. We deposited 3D DNA origami shapes, which were designed *in-silico* [38, 39] and folded in buffer containing MgCl_2_ (Figure 1a), on patterns of hydrophilic binding sites on hydrophobic substrates (Figure 1b). We adapted protocols described by Gopinath et al. for placing planar DNA origami on such patterned surfaces, which can be produced with e-beam lithography [1] or nanosphere lithography [36]. Effectively, both methods result in a hydrophobic surface with hydrophilic spots that act as the binding sites for the origami structures, we hypothesize primarily for the highly charged phosphate backbone of the DNA. We therefore employed two approaches to achieve upright placement of 3D DNA shapes: i) our hollow nanotubes, for example, have a small footprint in the desired upright position. In the undesired flat-lying orientation, in contrast, such a DNA tube exhibits a large contact area with the hydrophilic spots. Indeed, we observed mostly flat-lying tubes if they were administered directly onto the pre-patterned surfaces. We thus used a two-step process where planar DNA origami sheets were deposited first as connectors and the 3D DNA structures were annealed to these connector sheets in a subsequent step (Figure 1c). For this, specific anchor strands extend from the ends of the tubes and bind to strands protruding from the planar connector origami sheets. ii) Our barrels are designed such that the bottom-to-be faces of the structure are larger than its side and are therefore more likely to attach to the hydrophilic regions. Or, as in the case of the tetrapod, the four main faces are identical. These two objects were deposited directly on the patterned substrates (Figure 1d). After successful deposition, all samples can optionally be incubated with pre-hydrolyzed N-trimethoxysilylpropyl-N,N,N-trimethylammonium chloride (TMAPS) and tetraethylorthosilicat (TEOS) enabling the growth of a rigid silica shell to allow for drying of the products [8] (Figure 1e).

**Figure 1.**
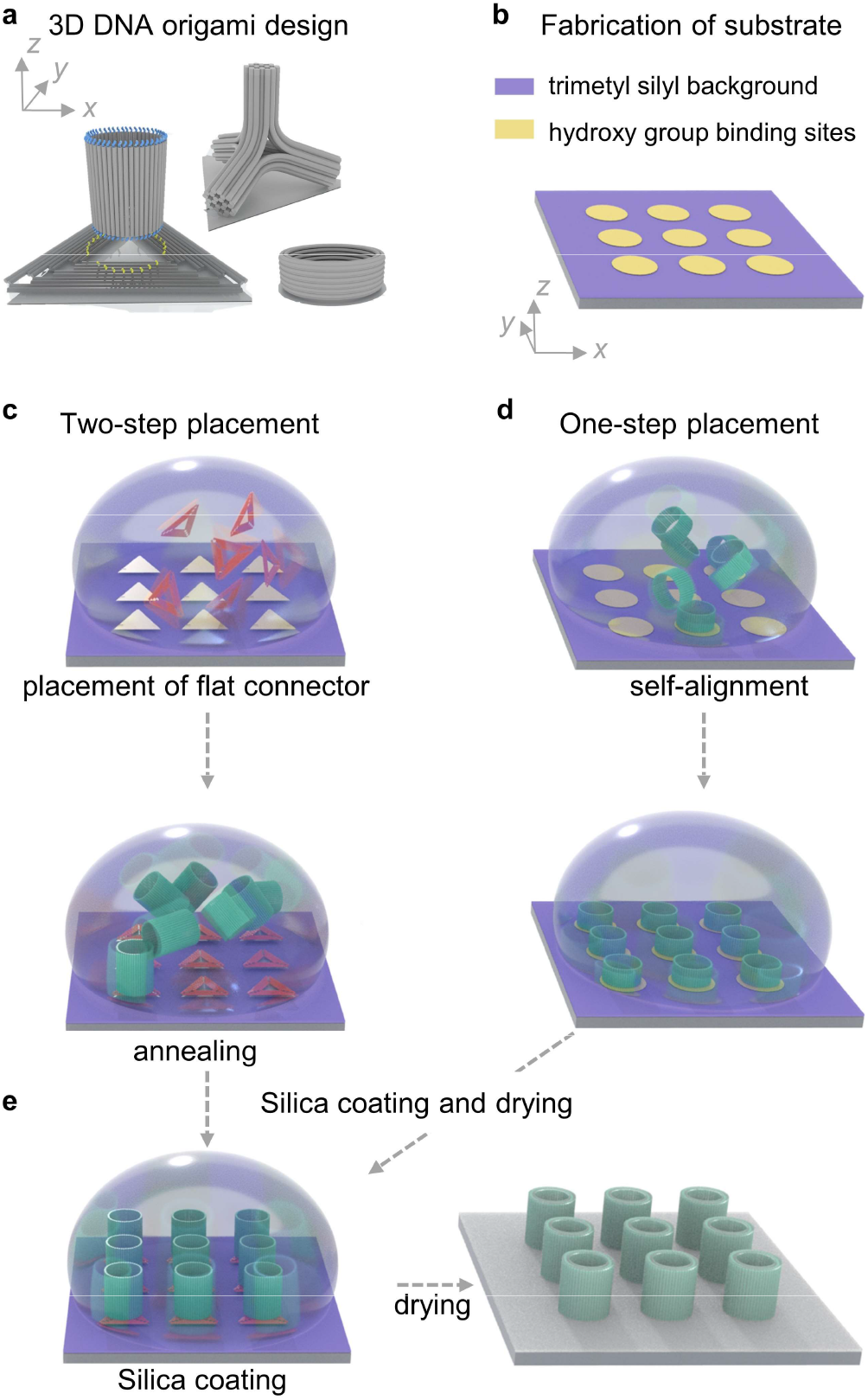
Assembly of 3D hybrid nanostructured substrates. a) Design of 3D DNA origami shapes and connection interfaces for on-surface assembly. b) Substrates are patterned by ebeam [3] or nanosphere lithography [36] to produce hydrophilic oxide patterns on HMDS-primed hydrophobic background (Si/SiO_2_ wafer or glass). c, d) Alignment and upright positioning of 3D DNA origami structures on patterned surfaces. DNA is represented in a cylinder model. Shapes that cannot self-align in an up-right position are placed in a 2-step process with planar DNA origami as connectors (Figure 1c). Other shapes are directly deposited to the patterned substrate (Figure 1d right). e) Growing rigid silica shells on the 3D DNA origami enables subsequent drying of the now rigidified objects.

The two-step placement method, which is based on sequence-specific DNA binding on a surface, enables us to position any 3D DNA origami shape in a defined directionality. As an example shape, we designed a DNA origami nanotube with a length of 50 nm and a diameter of 40 nm (Figure 2a). Here, a rolled-up single layer of 48 DNA duplexes forms the tube. Its native wall thickness is defined by the width of a DNA double helix, i.e. 2.1 nm (Supplementary Note 1, Figure S1 and Table S1). Figure 2a display a computer graphic of the tube and its built-in binding strands as well as the connector sheet. We used a variant of the “Rothemund triangle”. For one, this is a commonly used origami structure present in many laboratories and second, it has been positioned on lithographically patterned substrates successfully before [3]. Single-stranded DNA linkers extend from the center of the triangle roughly matching the circular footprint of the DNA tube [21]. Hybridization between these linkers and complementary anchor strands extending from the tube’s ends – we labeled both ends of the tubes with anchor strands to increase the probability of binding – brings the tubes and the triangles together in a way that the tubes “stand” on top of the triangles (Figure 2d, Supplementary Note 2, Figure S2). Figure 2b through 2d display transmission electron microscopy (TEM) images of the DNA objects where panel b shows a side view of a tube lying flat on the TEM grid, panel c the triangle and panel d the tube assembled on top of a triangle (additional TEM characterization in Figure S3 and Figure S4).

**Figure 2.**
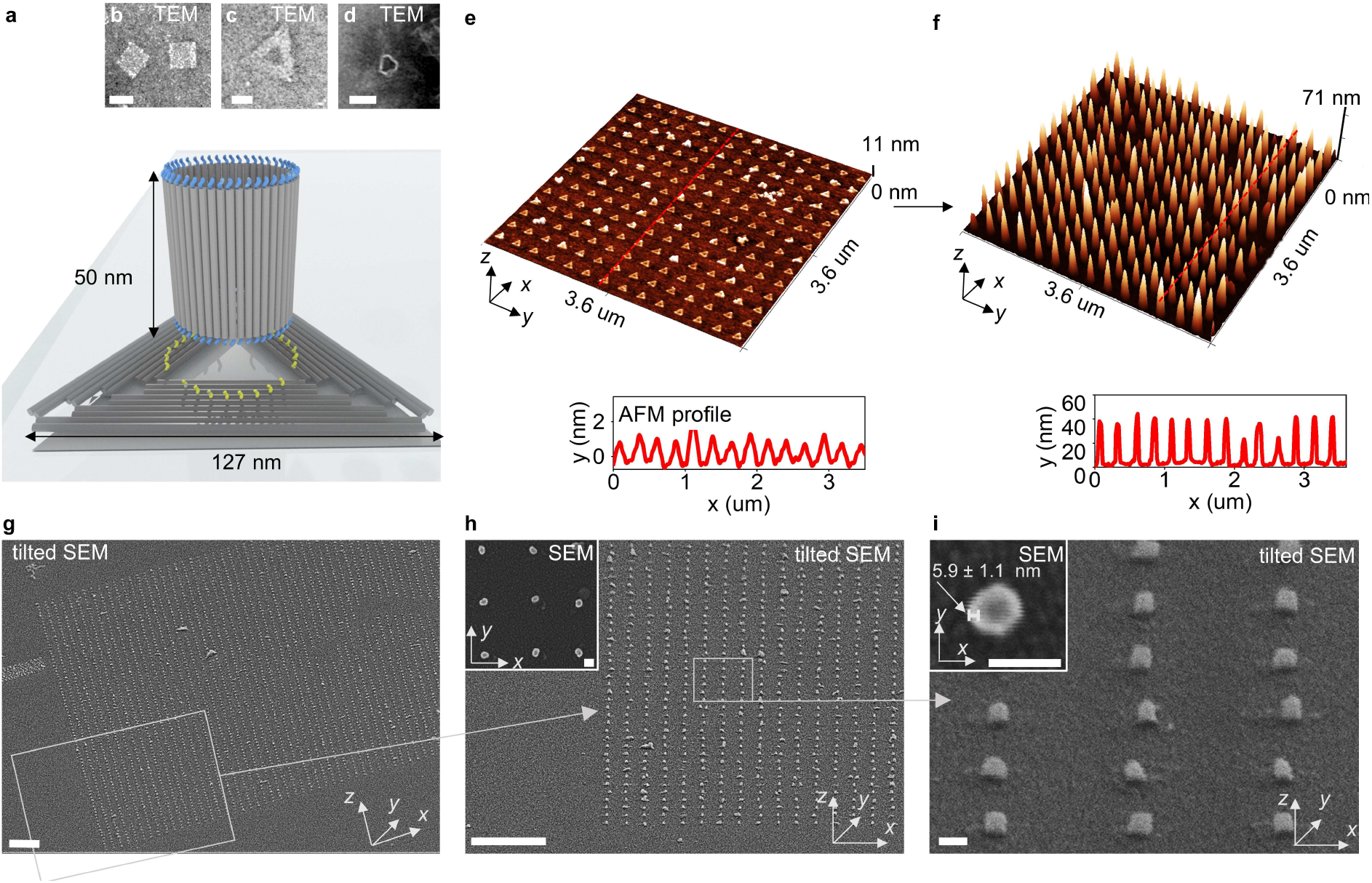
Assembly of 3D hybrid nanostructured substrates by on-surface annealing of DNA origami nanotubes to a flat connector origami. a) Design of the DNA origami tubes and triangles. Single-stranded DNA linkers extend from the center of the triangle roughly matching the circular footprint of the tube. Complementary anchor strands extend from the ends of the tubes. b-d) Uranyl-formate negative-stain TEM images of b) DNA tubes carrying 48 T_11_ ssDNA linkers, c) a triangle, carrying 27 A_12_ ssDNA anchors and d) a tube annealed with a triangle. e) AFM characterization of dried Si/SiO_2_ wafer with an array of DNA origami triangles carrying 27 A_12_ ssDNA anchors. f) AFM and g-i) SEM characterization of dried Si/SiO_2_ wafer with an array of silica-coated DNA tubes standing on top of triangles. Scale bars in g) and h): 1 μm. Scale bars in b – d, i) and in the inserts in h, i): 50 nm.

We fabricated Si/SiO_2_ wafers with square arrays of triangular binding sites situated 250 nm apart from each other *via* e-beam lithography. We deposited DNA triangles, carrying 27 ssDNA linkers each 12 nt in length, on a surface of patterned wafers. Generally, the mechanisms of origami-to-site binding are complex and small details in variations during the substrate fabrication can affect placement yields significantly [3]. The experimental conditions for high-quality positioning of planar origami on Si/SiO_2_ substrates (Tris buffer, pH of 8.35, Mg^2+^ concentration of 35 mM incubation time of 1h at RT, see Table S2 for all of the buffers used in the work) have been already reported in [3]. In order to reproduce the successful placement of triangles, we used these parameters and only adjusted the size of the binding sites and the DNA origami concentration (Supplementary Note 3). We achieved up to ~ 94% of sites occupied with a single triangle and ~ 5% of sites with multiple/aggregated triangles (Figures S5-S7).

Next, we incubated these pre-patterned surfaces with DNA tubes bearing 48 ssDNA anchors each 11 nt in length and complementary to the linker DNA on the triangles at 37°C for 1h in a Tris buffer with 12.5 mM MgCl_2_ (see Supplementary Note 4.1, Figures S8, S9 for details on in-solution assembly optimization and Supplementary Note 4.2, Figures S10-S13 for details on on-surface optimization). After annealing, the wafers with the full assemblies were exposed to a silica-coating procedure described in [8]. Subsequent air-drying and AFM and SEM imaging revealed rigid 3D DNA-silica nanotubes arranged in square arrays on the Si/SiO_2_ surface (Figure 2f). We observed up to 75 % occupancy of binding sites with individual standing tubes while the remaining 25 % of sites were doubly occupied (4 %), empty (2 %) or occupied with higher order aggregates (Figure S13b and Figure S14). The height of the silicified tube determined by AFM is ~ 45 nm, which is in good agreement with the designed tube length of ~ 50 nm. The wall thickness of the upright silica-DNA tubes is 6 ± 1 nm, as determined by SEM (insert in Figure 2i, Figure S15). This is below the state-of-the-art 10 nm resolution in 3D silica nanofabrication achievable by focused ion beam or thermal scanning probe lithography [40, 41].

Next, we studied how pattern diversity affects the placement and annealing yields. It was shown before that binding of triangles can be a function of array period, especially for binding sites located towards the center of a given array. Occupied sites may inhibit the 2D diffusion of unbound DNA objects to unoccupied sites and so the occupation rate decreases as the period decreases [3]. We examined triangles binding to sites in square arrays with periods ranging between 170 nm and 400 nm on the same Si/SiO_2_ chip. To our delight, we did not observe the expected drop of binding rates with a decrease of period from 400 nm to 170 nm (Figure 3a,c, Figure S16). In fact the highest percentage of 95% of sites binding a single triangle for the 250 nm period is slightly decreased to 93% for the 400 nm period and to 91% for the 170 nm period. Consequently, the site occupancy and alignment of standing tubes in arrays of corresponding periods did not vary significantly. We achieved up to 71% and 74% of sites with individual upright nanotubes for the 170 nm and 400 nm arrays, respectively (Figure 3, b-d, f-h, Supplementary note 5, Figure S17). This opens a route to create diverse patterns and arrays of integrated 3D DNA-silica nanodevices on one chip.

**Figure 3.**
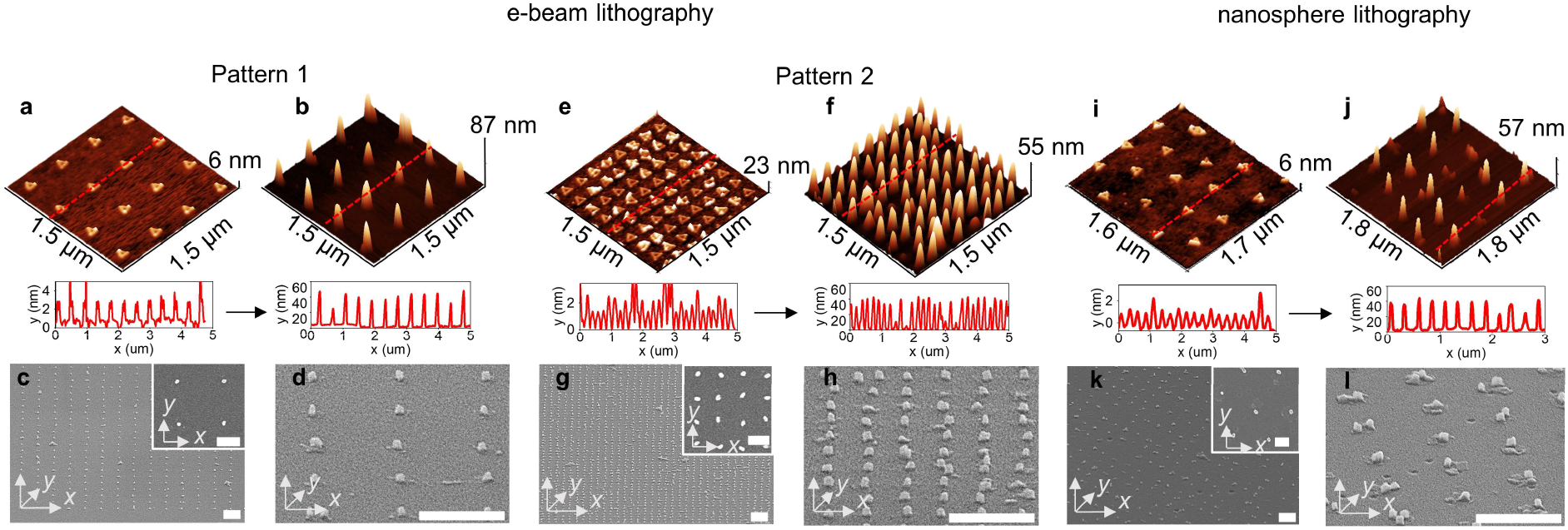
Pattern diversity. a-h) AFM and SEM characterization of dried Si/SiO_2_ surfaces with square arrays of DNA origami prepared with e-beam lithography with periods of 400 nm (a-d) and 170 nm period (e-h). a, e) AFM characterization of surfaces with square arrays of DNA origami triangles carrying ssDNA linkers. b, f) AFM and c, d, g, h) SEM characterization of surfaces with arrays of silica-coated DNA tubes standing upright on the triangles. i-l) AFM and SEM characterization of a dried glass surface with a hexagonal pattern of DNA origami prepared *via* nanosphere lithography [36]. i) AFM characterization of the glass surface with a hexagonal array of DNA origami triangles carrying ssDNA linkers. j) AFM and k, l) SEM characterization of dried glass chip with a hexagonal array of silica-coated DNA tubes standing on top of triangles. c, d, g, h, k, l) are tilted SEM images, inserts are top-view SEM images. All scale bars: 400 nm. Scale bars in the inserts in c, g, k): 200 nm.

For many applications it can be advantageous to use a cleanroom-free, large-scale DNA origami placement method such as the benchtop technique of nanosphere-DNA origami lithography described in [36]. In nanosphere lithography, a layer of close-packed nanospheres creates a crystalline pattern of contact points for the selective passivation of the supporting glass substrate [36]. After chemically rendering the “free” glass surface hydrophobic, the nanospheres are lifted off and DNA origami structures can subsequently bind to the hydrophilic, close-packed, hexagonal pattern defined by the previous contact points.

We utilized this method of bottom-up nanopatterning for the two-step placement described above. Although nanosphere lithography creates circular binding sites, we achieved similar success in triangle placement as Shetty et al. [36] did with circular DNA origami (Figure 3i, Figure S18). The yield of single tubes attaching to the triangles increased with the incubation time (Figure S19), but it never came close to the values obtained for e-beam lithography patterning on Si/SiO_2_ wafers. After 3 h of annealing at 37°C, the occupancy of sites reached ~ 78 % with only ~ 35 % of sites carrying a single standing tube. Further incubation (up to 24 h) leads to almost full occupancy of the binding sites (~ 97 %), however, with still only ~ 36 % of sites with single tubes accompanied by a dramatic increase in multiple binding events. This could be a result of the geometrical mismatch between the circular biding sites and the triangular DNA origami connectors, which leaves free hydrophilic binding space where the DNA tubes can bind directly. Noteworthy, our hexagonal arrays of sphere-lithography assembled DNA tubes cover a total area of more than 4 mm^2^ (Figure S20), proving that it is possible to arrange complex 3D DNA-silicified molecular breadboards over macroscopic areas in a conventional wet lab environment.

Another approach to binding the 3D origami shapes to substrates relies on direct binding and self-aligning. To achieve this, it is necessary to either design the objects such that certain faces of are more likely to attach to the hydrophilic binding sites or that all main faces are identical.

As an example of the first case, we demonstrate up-right positioning of DNA origami barrels. Our DNA origami barrel is a donut-shaped structure designed previously [9] and constructed from horizontally aligned, circular DNA duplexes. The barrel has a diameter of 60 nm and a height of 27 nm (Figure 5a-c, Figure S21). The relatively thick walls and a low aspect ratio of 1:2.2 (height to width) promote the horizontal binding of the structures. Moreover, the binding sites were tuned to match the diameter of the barrels. We fabricated a series of Si/SiO_2_ wafers *via* e-beam lithography with circular binding sites, varying the diameter between 75 % and 200 % of the actual barrel diameter of 60 nm. We obtained the best yield of single bound upright barrels (70 %) with spot sizes of 45 nm while the percentage of the barrels lying on side is reduced to 16 % (Figure 4d,e and Supplementary Note 6, Figures S22-S26).

**Figure 4.**
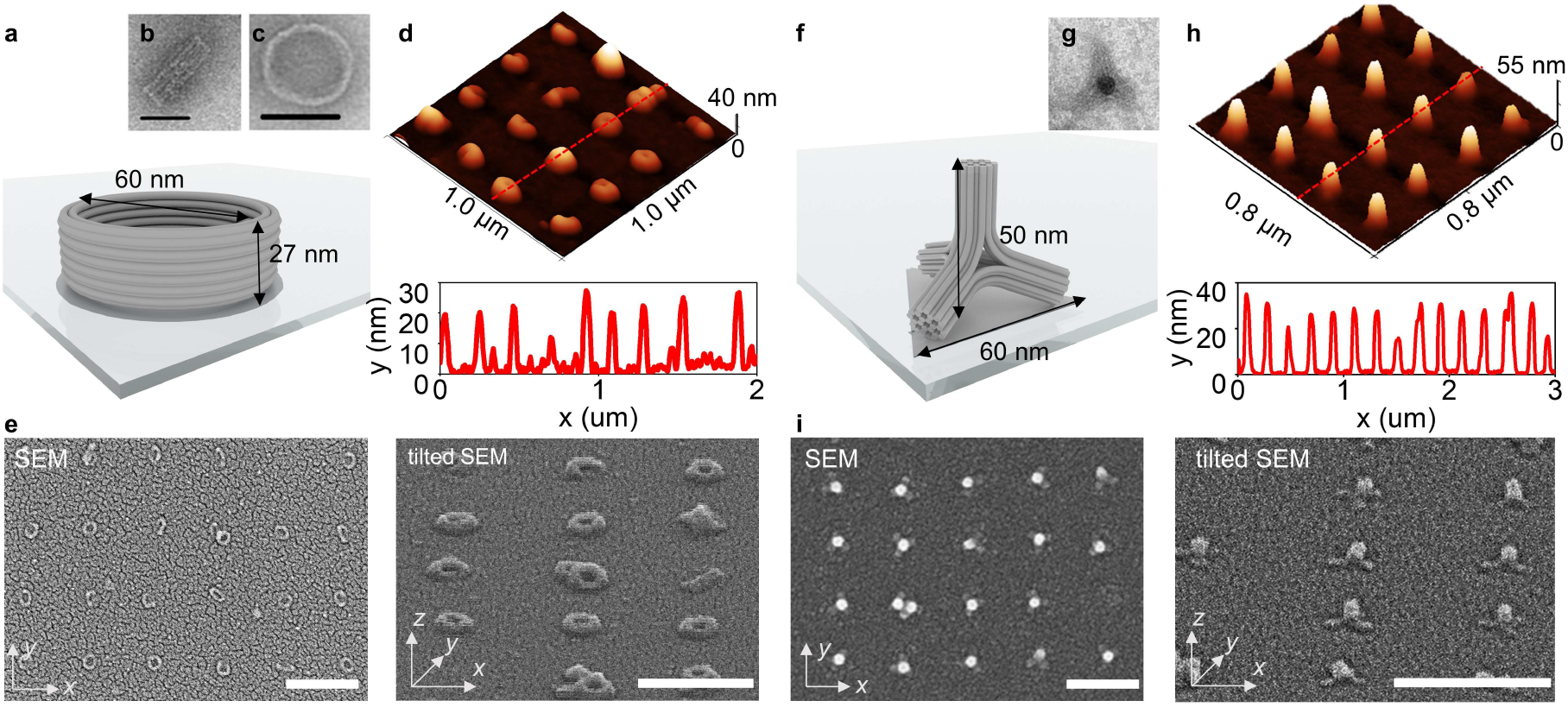
Assembly of 3D hybrid nanostructured substrates by direct deposition a) Design of the DNA origami barrels, illustrated as a cylinder model. b, c) Uranyl-formate negative-stain TEM images of b) a DNA origami barrel lying on its side, c) an up-right DNA origami barrel. d, e) AFM and SEM characterization of a dried Si/SiO_2_ substrate with a square array of silicified DNA origami barrels. f) Design of the DNA origami tetrapods. g) Uranyl-formate negative-stain TEM images of the DNA origami tetrapod. d, e) AFM and SEM characterization of a dried Si/SiO_2_ substrate with a square array of silicified DNA origami tetrapods. Scale bars in e, i): 400 nm.

**Figure 5.**
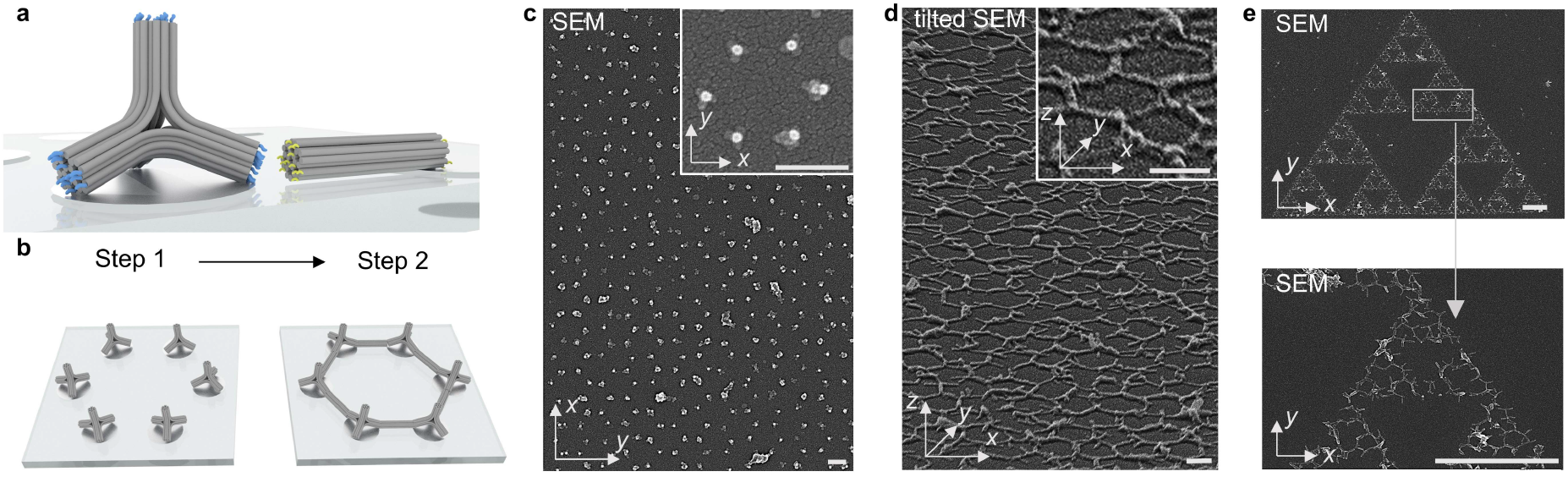
Assembly of 3D hybrid periodic networks on substrates by on-surface annealing of DNA origami 24HBs to the tetrapods, pre-absorbed on the binding sites. a) Design of the interface between the DNA origami tetrapods and the 24HBs. 12 single-stranded DNA linkers extend from each leg of the therapod. 12 complementary anchor strands extend from each side of 24HB. b) Assembly of the periodic networks in a two-step process. First, DNA origami tetrapods absorb to the binding sites arranged in a honeycomb lattice (Step 1). 24HBs connect neighboring tetrapods pairs in an annealing process (Step 2). c) SEM images of Si/SiO2 substrate in a honeycomb lattice of tetrapods bearing 12 ss-DNA linkers. d-e) SEM characterization of the same sample as in (c), interconnected with 24HB bearing 12 anchor strands from both sides. The resulting continuous network covers a micron-sized honeycomb pattern (d) and can be designed as a fractal arrangement of individual hexagons forming Zelinski triangle (e). Scale bars in c, d): 200 nm. Scale bars in e): 2 μm.

As an example of the second design case, we employed a DNA origami tetrapod that consists of four equivalent arms, resulting in a four-fold symmetric object (Figure 4f, g and Figures S27, S28). Each arm is composed of interconnected 24-helix bundles (24HBs) as described in Supplementary Note 7. As expected, after incubating the origami tetrapods on patterned substrates, we observed individual objects standing on three legs on the binding sites (in Figure 5f). After optimizing binding site shape and size, we achieved occupancy yields of 79 % while single therapods occupied 43 % of the sites and 36 % of the sites carried multiple tetrapods (Supplementary note 8, Figures S29-S32). Square arrays with a 200 nm period of silica coated and dried tetrapods imaged by SEM and AFM are presented in Figure 4h and i, respectively. The height of an individual silica-coated tetrapod obtained from AFM measurements of dried samples is ~ 40 nm, which is in fair agreement with the designed 50 nm. Most likely, the tetrapod “sits” flat on the surface, reducing the effective height of the structures. We assume that a similar approach can be used for the individual placement of all 3D shapes with equilateral faces such as tetrahedrons, cubes, octahedrons, etc. Overall, while the arrangement of deposited 3D structures in this one-step method is worse compared to the two-step process, it can be helpful for positioning a large variety of 3D shapes with low-aspect ratio as cuboids, cylinders, cones etc.

### Further assembly

In principle, individually placed 3D DNA shapes can be used as seeds for further assembly in *xyz* directions in subsequent surface annealing steps. Both the DNA shapes and the lithographic pattern can be rationally designed to create not just periodic arrays but complex 3D networks. As a proof-of-concept, we created large-scale periodic honeycomb networks by a 2-step placement procedure with the initial placement of tetrapods followed by the annealing of 24 HBs to the legs of the tetrapods (Figure 5a, b). Here, tetrapods bearing 12 ss-DNA linkers on each leg were deposited on lithography-patterned surfaces as described above and arranged with a spacing adjusted to accommodate a 24HB with 12 complementary anchor strands extending from both sides (Figure 6c, Supplementary notes 9 and Figures S33, S34). The resulting 3D network is designed as a micron-sized honeycomb array (Figure 6d) or as a fractal arrangement of individual honeycombs forming Zelinski triangles (Figure 6f).

## SUMMARY

In this work we advanced from 2D to 3D in the placement of individual origami objects on lithographically defined surfaces. In addition to avoiding multiple binding per sites and empty binding sites, we achieved full spatial control over a deposition of a given 3D shape on a certain face. We prevented 3D DNA origami structures from collapsing upon drying by coating the DNA with silica [8]. Moreover, and importantly for future applications of such DNA-assembled 3D substrates, nanometer-precise modification with a wide variety of nanoscale components can be directly implemented on these DNA origami shapes. Individual placement of 3D DNA origami on macroscopic patterns could potentially boost conventional methods of lithographic shaping of diverse substrates and organizing matter at nanoscale for future ultra-scaled 3D manufacturing. We envision that such strategies may bring a high degree of design freedom and versatility into nanofabrication.

## DETAILS ON AUTHORS CONTRIBUTIONS

T.L. and I.M. designed this study. I.M., E.E. and V.R. designed, assembled, purified DNA origami samples, designed and optimized interfaces. G.P. and X.Y. designed, assembled, purified DNA origami tetrapods and 24HBs and designed the tetrapod-24HB interface. I.M., E.E. V.R. and M.D. performed placement experiments, surface annealing experiments, AFM and SEM measurements and data analysis with the assistance from P.A.. I.M. and T.L. wrote the manuscript with input from all authors.

## ACKNOWLEDGMENTS

We thank Christian Obermayer for clean room assistance and Susanne Kempter for assistance with TEM. We thank all members of Tim Liedl Group for and for helpful discussions. I.M., E.E., G.P. and T.L. acknowledge funding from the ERC consolidator grant “DNA Funs” (Project ID: 818635) V.R., X.Y. and T.L. further acknowledge support from the cluster of excellence e-conversion EXC 2089/1-390776260.

## ONLINE CONTENT

Materials and methods, supplementary information, additional references, can be found online

