## Supplementary Information for "Site-directed placement of three-dimensional DNA origami"

**Geschwister-Scholl-Platz 1, 80539 Munich, Germany**

Contents

Supplementary Notes 1-9

Supplementary Figures 1-34

Supplementary Tables 1, 2

Materials and Methods

Additional References

### Supplementary Note 1. Design of the DNA origami nanotube

DNA origami nanotube is a hollow cylinder consisting of a single layer of 48 double-stranded DNA duplexes aligned side-by-side and held together with crossovers, as shown in Supplementary Information Figure 1. The parameters of the nanotubes (number of duplexes $N_d$, dihedral angle, intrinsic curvature, diameter) are listed in the Supplementary Table 1. The positions of crossovers were calculated using the computational algorithm The NanoCooper [1]. The design was entered manually into Cadnano v2 [2] and further corrected in order to avoid mechanical isomerization between two possible conformations of DNA nanotubes [3]. Previously, it was found that tubular designs of origami structures can have two stable conformations: a short and a long tubular form that may stem from different isomers of Holliday junctions contained inside the origami [4, 5]. Mechanical isomerization is only possible in DNA origami with rotational symmetric arrangements of Holliday junctions [3]. Therefore, we shifted the position of crossovers in order to break the rotational symmetry of Holiday junctions in the nanotubes.

### Supplementary Note 2. Design of the interface between the nanotube and the triangle

For the hybridization of nanotubes with DNA origami triangles, 24 staples on each edge of the tube were modified by extending 11 thymine nucleotides on the 3′ end (Supplementary Figure 2). These T11-modified DNA staples were introduced into the DNA scaffolds in place of the original DNA staples and folded and purified as described in 0.

The 'sameside sharp triangle' design is taken from [6]. The "sameside triangle" has all nick positions on the same face of the triangle so that ~200 modifications can be made to the same side of the triangle. We further modified 27 staples close to the middle hole by extending with 8, 12 or 20 adenine nucleotides on the 5′ end (see Supplementary Figure 2). Since it cannot be controlled or predicted which side of a sameside triangle will be facing the patterned surface, we further positioned the nicks of corresponding 27 extended staples in the plane of the triangle and pointed them towards the triangular hole in the middle. Therefore, extended strands can point out towards each side of the triangle with a 50% probability.

9 of these extended staples lie on the two DNA helices closest to the triangular hole in the center of the triangle and form a circle with a diameter of ~32 nm, as shown in Supplementary Figure 2. These extensions strands, labelled as "red" extensions in the following, are expected to have a higher flexibility for changing from extending from one side of the triangle to the other due to their closeness to the central hole of the triangle.

The remaining 18 extensions strands lie on DNA helices further away from the center and form a larger circle of ~44 nm, Supplementary Figure 2. They are labelled as "violet" extensions. Diameters of both circles are close to the designed ~40 nm diameter of nanotube.

These A-modified DNA staples were introduced into the DNA scaffolds in place of the original DNA staples as three different sets: only red extensions (small circle), only violet extensions (large circle) or both (all 27 extensions at once).

Additionally, the number of adenine nucleotides (8, 12 or 20) was varied for each set of extensions, providing wide variability in length, number and geometry of extensions for the precise optimization of the tube-triangle hybridization interface. Triangles with all 9 possible modification sets were assembled under same folding and purification conditions as described below in Materials and Methods.

### Supplementary Note 3. Optimization of DNA origami triangles placement on Si/$SiO_2$

We were able to reproduce successful placement of triangles using the substrate preparation protocol and placement buffer parameters reported previously in [6] from the first attempt. We further adjusted only the binding site size via constructing arrays with binding sites of varying sizes, from 70% to 100% of a triangular origami edge length (127 nm). Besides, we varied the dosage of e-beam exposure (400 uCu/cm$^2$, 500 uCu/cm$^2$, 600 uCu/cm$^2$), because it also influences the final size and shape of the binding sites. Additionally, we adjusted the concentration of DNA origami (100 pM, 200 pM, 300 pM). After placement, wafers with triangles were dried and imaged with dry mode AFM. Each AFM image was processed using the Gwydion software. We measured the binding site occupancy (percentage of sites with one or more origami), number of origami at a site (1 or multiple/aggregated) by hand-annotating images as shown in Supplementary Figure 5.

The optimum parameters we found for DNA origami placement are triangular binding site with 120 nm edges, 400 uCu/cm$^2$ dosage of e-beam exposure and DNA origami concertation of 300 pM. A typical AFM image of a wafer with triangles arranged in square lattice with 250 nm period is presented in Supplementary Figure 6. For these optimal placement parameters, the number of sites was analyzed in 3 independent replications and a histogram with binding statistics is presented in Supplementary Figure 7. The site occupancy was up to 99% and single triangle binding was up to 95%.

### Supplementary Note 4. Optimization of nanotube annealing with triangles on Si/$SiO_2$

#### *4.1 Optimization in Solution*

To push nanotube-triangle binding yields towards 100%, we first predetermined optimal parameters (number, length and spatial geometry of DNA linkers) in solution experiments.

DNA Origami nanotubes with 11T extensions and DNA Origami triangles with different sets of polyA extensions, were folded and purified as described in Materials and Methods.

Three different sets of triangles extensions were used (Supplementary Figure 2): red set extensions (R) with 9 polyA strands arranged in a small circle with a diameter of approximately 32 nm close to the triangular hole in the center of the triangle, violet set extensions (V) with 18 polyA strands arranged in a larger circle with a diameter of approximately 44 nm further away from the triangular hole in the center of the triangle, or both sets of extensions simultaneously (RV) with 27 polyA strands. Additionally, each set of extension has 3 variations in number of bp of 8A, 12A and 20A.

Nanotubes and triangles with all possible variants of extensions (R20A, V20A, RV20A, R12A, V12A, RV12A, R8A, V8A, RV8A) were mixed at a 1:1 ratio and annealed in Folding buffer. Annealing of the nanotubes with triangles results in the formation of long "bamboo" structures, consisting of alternating triangles and nanotubes hybridized to one another. The length and accuracy of these bamboo structures is an indicator for the quality of the hybridization and is evaluated by gel electrophoresis and TEM (Supplementary Figure 8 and Supplementary Figure 9, respectively).

Triangles with RV 12A extensions were selected for further experiments.

Additionally, we investigated the influence of $MgCl_2$ concentration (10 mM – 40 mM) and incubation time (1 h – 4 h) on the annealing yield of 11T nanotubes and RV 12A triangles in the Placement buffer. No vast differences in annealing yields in studied at the syudied $MgCl_2$ concentration and time ranges were revealed by gel electrophoresis and TEM (data not shown).

#### *4.2 Optimization on the surface of Si/SiO$_2$ wafers*

We studied the influence of the number and geometry of polyA extensions on triangles, concentration of nanotubes and $MgCl_2$ concertation in a Hybridisation buffer on the yield of single nanotube attachment to a triangle which is pre-absorbed on a binding site on a Si/SiO$_2$ wafers.

For this, we prepared Si/SiO$_2$ wafers with 120 nm triangular binding sites arranged in arrays with 250 nm period and placed with DNA origami triangles. Then we introduced nanotubes bearing 48 11T extensions on both ends in Hybridization buffer to the surface of the wafers and annealed them to the pre-absorbed triangles as described in Materials and Methods.

In order to study the influence of number and geometry of extensions on triangle and nanotube binding yield, we used triangles modified with 18 12A extensions (V12 triangles) and 27 12A extensions (RV12 triangles), arranged as shown in Supplementary Figure 2. Additionally, we adjusted the concentration of nanotubes (250 pM, 500 pM) and concentration of $MgCl_2$ (35 mM, 12.5 mM) in the Hybridisation buffer. After surface annealing, wafers with nanotubes bound to triangles were silica-coated, dried and imaged with SEM. Then we calculated the statistics of single standing tube binding. We evaluated the percentage of single standing tubes at a site (1 or 2 standing nanotubes), nanotube alignment (percentage of nanotubes lying on side/aggregated) and binding site occupancy (percentage of sites without origami), by hand-annotating SEM images as shown in Supplementary Figure 10.

Binding statistics and SEM images of dried silica-coated nanotube arrays as a function of number and geometry of 12A extensions on triangles, nanotube and $MgCl_2$ concertation in Hybridisation buffer are shown in Supplementary Figure 11, Supplementary Figure 12 and Supplementary Figure 13, respectively.

Optimum parameters chosen are RV12 triangles (Supplementary Figure 11), 250 pM nanotube concentration (Supplementary Figure 12) and 12.5 mM $MgCl_2$ concentration (Supplementary Figure 13). For these optimal placement parameters, the number of sites was analyzed in 2 independent replications. A histogram with binding statistics is presented in Supplementary Figure 13. The maximum achieved site occupancy with single standing tubes was 75%.

### Supplementary Note 5. Nanotube-triangle annealing on arrays with different periods on Si/SiO$_2$

In order to prove that nanotube-triangle annealing yield is independent on the nanopattern, square arrays of binding sites with three different periods (170 nm, 250 nm, 400 nm) were fabricated on the same Si/SiO$_2$ chip.

Different periods of 170 nm, 250 nm, 400 nm between triangles result in comparable single triangle binding yields of 91.00%, 95.39%, 93.06%, respectively. AFM images of dried Si/SiO$_2$ wafers are presented in Supplementary Figure 16.

Consequently, the yield of single tube binding is almost the same for the different periods (71% for 170 nm, 75% for 250 nm, 74% for 400 nm), as shown in Supplementary Figure 17.

### Supplementary Note 6. Optimization of DNA origami barrels placement on Si/SiO$_2$

Si/SiO$_2$ wafers were nanopatterned as described in Materials and methods and the purified barrels are placed on the surface using placement conditions optimized previously for flat triangular DNA origami: TE placement buffer with pH 8.35, 35 mM MgCl2 and incubation time of 1 h [10]. The origami is placed on circular binding-sites with a diameter of 55 nm defined with an e-beam exposure of 400 uC/cm$^2$. The SEM image of a wafer with patterned barrels with a concentration of 200 pM is shown in Supplementary Figure 22. It is observed that the placement protocol itself works as almost all binding sites are occupied and the patterned ring structures represent standing barrels. Apart from standing barrels, the patterned area contains single barrels lying sideways, multiple binding of aggregated origamis on one binding site and empty binding sites without barrels. Therefore, the parameters of the placement were further optimized. The goal is to observe the changes in placement yield for different conditions and ultimately to find the optimal parameters for the highest percentage of placement of single standing barrels. Supplementary Figure 22 depicts examples for the considered binding events. The standing barrel (green) is circle-shaped with a clearly visible hole in the middle. A binding event is considered as a standing barrel when the hole is clear and in contrast to the structure. Lying barrels (yellow) are also counted as single binding when they are the only structure on a binding-site. Multiple binding is shown in blue. Three barrels are located on one site, the holes in the structures are visible. It is also counted as a multiple binding event. An empty binding site is marked in red. No structure is bound to the surface, only the background is visible.

After optimizing of origami concentration (Supplementary Figure 23), exposure dosage (Supplementary Figure 24) and binding-site size (Supplementary Figure 25), the best yield of single standing barrel binding was achieved for the following parameters: 100 pM concentration, 400 mC/cm$^2$ dosage and a binding-site size of 45 nm (75% of the actual DNA origami barrel diameter). Supplementary Figure 26 shows a SEM image of the optimal placement achieved. Using these experimental conditions, it is possible to obtain a binding-site occupancy of 97.96%. The amount of standing single binding events is 70.41% and only 12.25% of sites have multiple binding.

### Supplementary Note 7. Design of DNA Origami Tetrapod

The DNA origami tetrapods were designed in caDNAno [2] as shown in Supplementary Figure 27.

They are composed of four 35 nm long “arms” with a round 24HB cross-section and 15 nm diameter which are designed to be oriented at a 109.5° angle with respect to each other.

### Supplementary Note 8. Optimization of DNA origami tetrapod placement on Si/$SiO_2$

Si/$SiO_2$ wafers were nanopatterned and the purified tetrapods are placed on the surface as using placement conditions: TE placement buffer with pH 8.35, 35 mM $MgCl_2$ and incubation time of 1 h [10]. The origami is placed on square binding-sites with a diameter of 35 nm defined with an e-beam exposure of 400 mC/cm$^2$. The SEM image of a wafer with patterned tetrapods with concentration of 400 pM is shown in Supplementary Figure 29. The SEM image depicts examples for the considered binding events. The standing tetrapod (green) is has clearly visible three legs and fourths leg points towards the image plane. Lying or deformed tetrapods are marked by blue, as well as multiple binding or aggregates. An empty binding site is marked in red.

After optimizing exposure dosage (Supplementary Figure 24) and binding-site size (Supplementary Figure 25), it was concluded that overall the best yield of single binding was achieved for the following parameters: 400 pM concentration, 600 mC/cm$^2$ dosage and a binding-site size of 35 nm. Supplementary Figure 32 shows a SEM image of the optimal placement achieved. Using these experimental conditions, it is possible to obtain a binding-site occupancy of 81%. The amount of standing single binding events is 44%, 37% of sites have multiple binding and 19% are empty binding sites.

**Supplementary Note 9. Design of the interface between the tetrapod and the 24HB**

The idea is to use the tetrapod as the placement base on patterned $Si/SiO_2$ surfaces and interconnect every pair of neighbouring tetrapods with a 24HB.

First, we designed 24HB extension in caDNAno [2] as shown in Supplementary Figure 33. The design is based on the p7249 scaffold which yields a bundle with 105 nm length and 15 nm diameter.

Next, we extended 12 staples on each surface of a tetrapod with the sequence GGGAAGGG (Supplementary Figure 34). The 24hb extensions have 12 anchors with complementary sequences arranged in the same pattern on each end surface.

Supplementary Figure 1: DNA Origami Nanotube Design

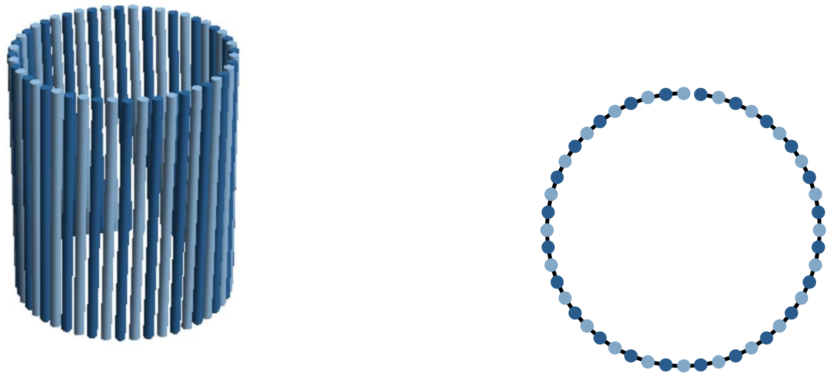

Schematic of nanotube constructed from 48 double-stranded DNA duplexes aligned side-by-side and held together with crossovers. DNA duplexes shown as light blue and dark blue cylinders with rightward and leftward polarity respectively. Side (left) and along-axis view (right) of DNA origami nanotube. Schematic of the nanotube was created by the computational algorithm The NanoCooper [1] with nanotube design parameters listed in Supplementary Table 1.

Supplementary Figure 2: Design of the interface between the nanotube and the triangle

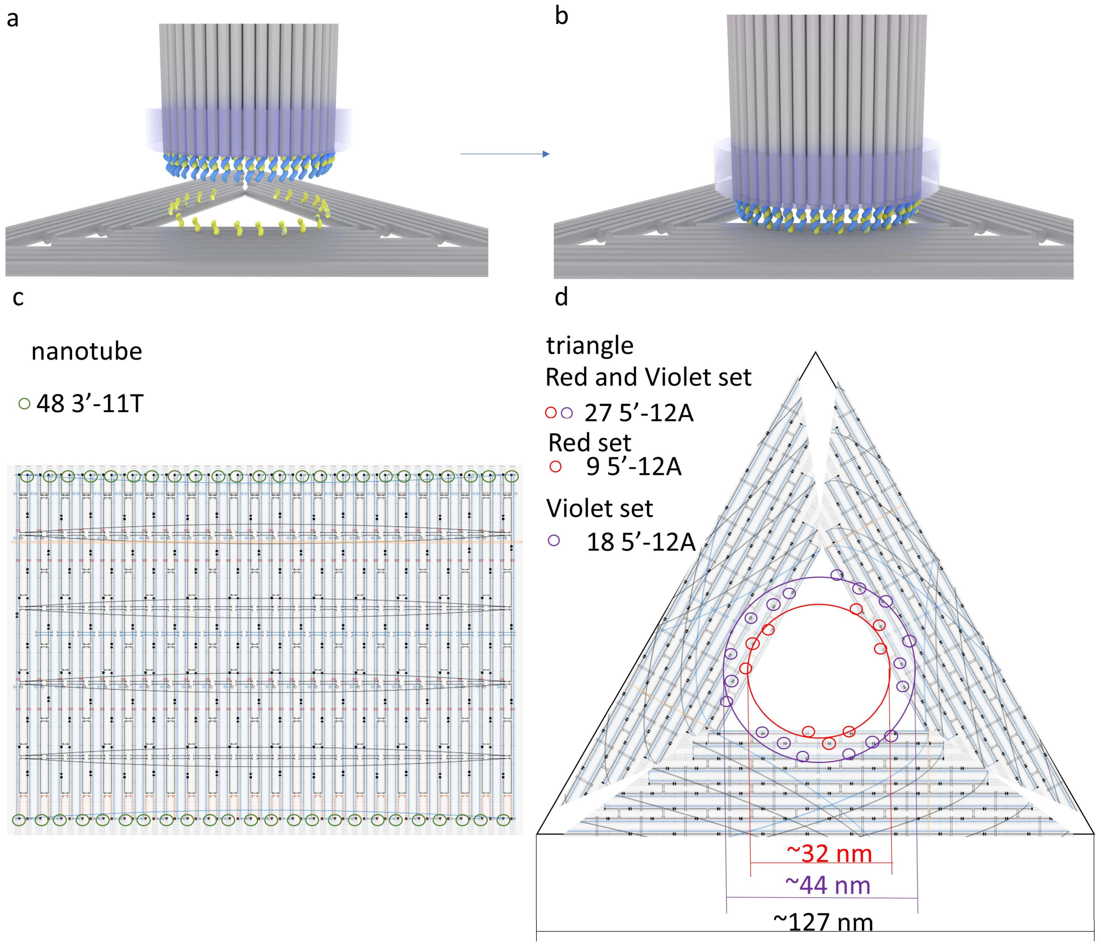


Design of the interface between nanotubes and triangles. (a). Single-stranded DNA linkers extend from the center of the triangle roughly matching the circular footprint of the nanotube. Hybridization between these linkers and complementary anchor strands extending from the nanotubes' ends brings the tubes and the triangles together so that the tubes "stand" on top of the triangles (b). (c) Green circles indicate 48 staples on both edges of the nanotube that were modified by extending 11 thymine nucleotides on the 3′ ends. (d) Red and violet circles indicate the 27 staples on the triangle which are extended by 8, 12 or 20 adenine nucleotides on the 5′ end. The staple diagrams of the nanotube (left) and triangle (right) were exported from caDNAno, the poly(A)- or poly(T)- functionalized positions were added by hand.

Supplementary Figure 3: TEM image of the nanotubes

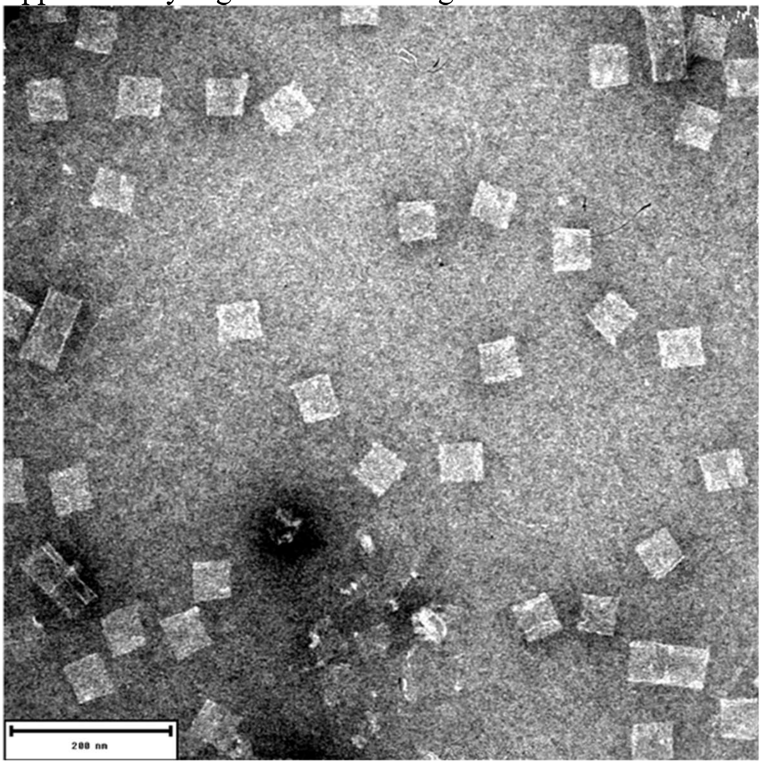


TEM image of DNA origami nanotubes with PolyT extensions (11T) at both ends after Amicon purification. DNA Origami tubes are lying on the side and are flattened on the TEM grids. Scalebar: 200 nm.

Supplementary Figure 4: Size distribution of nanotubes

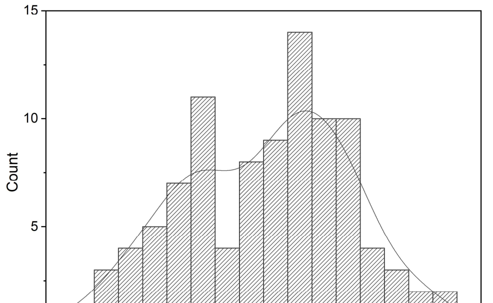


Nanotube size distribution from SEM. The length of the nanotube is approximately ~48 nm (left peak), the half of the circumference is ~53 nm (right peak). The calculated diameter of the nanotube is ~34 nm.

Supplementary Figure 5: Data analysis of dried DNA origami triangle arrays on Si/$SiO_2$ wafers

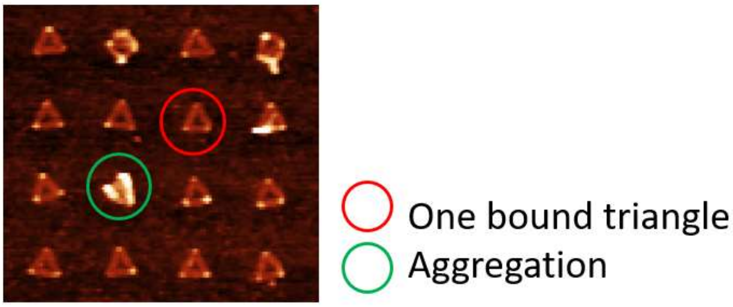


Dry-mode AFM images of Si/$SiO_2$ wafers with triangles bearing 27 12A extensions arranged in a square lattice with a period of 250 nm. Examples of binding events are marked: single binding (green), multiple binding/aggregated (red).

Supplementary Figure 6: AFM images of dried DNA origami triangle array on Si/$SiO_2$ wafers

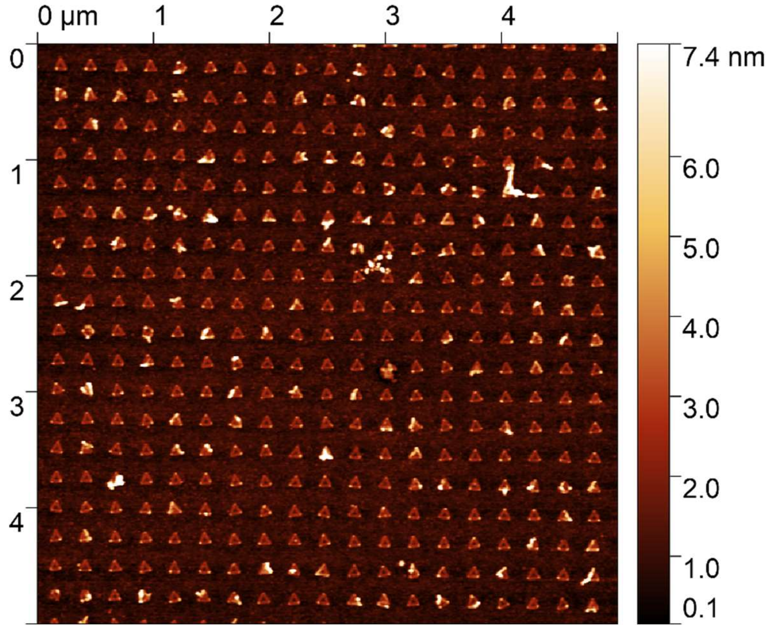


Dry-mode AFM images of Si/$SiO_2$ wafers patterned with triangles bearing 27 12A extensions with 250 nm period between triangles.

Supplementary Figure 7: Histograms with placement statistics of DNA origami triangle arrays on $Si/SiO_2$ wafers with 250 nm period and 120 nm binding site size.

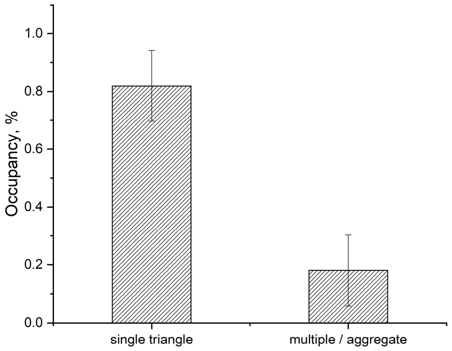


This histogram shows the quality of triangular origami placement in arrays on Si/SiO2 wafers with 250 nm period and 120 nm triangular binding site size, 400 $uCu/cm^2$ and 300 pM DNA origami concentration. Error bars are SD for N = 3 independent replications (of placement, washing, etc.) using different chips from the same wafer. 361 binding sites were scored for each replicate.

Supplementary Figure 8: Ooptimization of number, length and geometry of PolyA extensions on triangles

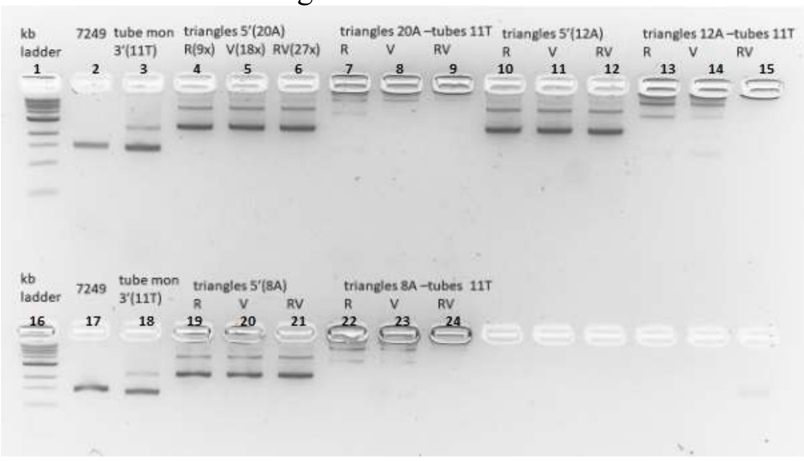


Gel Electrophoresis of nanotubes modified with 48 11T extensions from both sides (lines 3, 18) and triangles with different polyA extensions sets R (lines 4, 10, 19), V (lines 5, 11, 20) and RV (lines 6, 12, 21) as shown in Supplementary Figure 2 and different extension lengths 20A (lines 4-6), 12A (lines 10-12) and 8A (lines 19-21). 11T nanotubes and each type of triangles were mixed together at a concentration of 5 nM and incubated at 37°C for 1 hour in a Folding buffer with 12.5 mM $MgCl_2$ and loaded to the agarose gel immediately after the incubation. Conditions which we identified as optimal were triangle 12A – nanotube 11T and triangle 8A – nanotube 11T in RV configuration (lines 15 and 24, respectively). These samples were imaged under TEM (Supplementary Figure 9).

Supplementary Figure 9: TEM images of nanotubes and triangles with 8A or 12A extensions annealed in the buffer

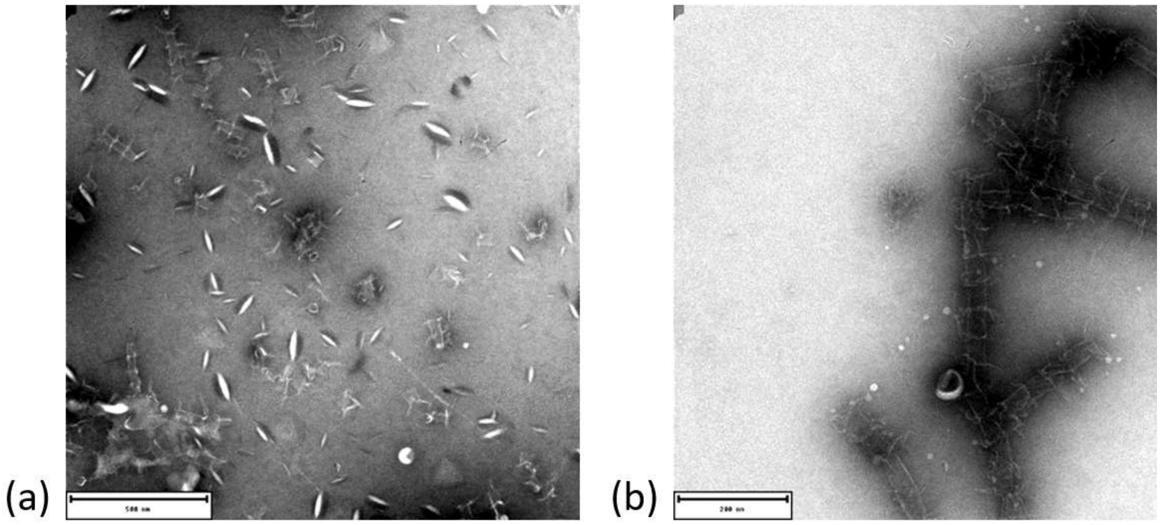


TEM images of 11T tubes and triangles with different types of extensions, mixed 1:1 at 5 nM, in Folding buffer with 12.5 mM $MgCl_2$, incubated for 1h at 37° a) RV 8A triangles and 11T tubes, b) RV 12A triangles and 11T tubes: bamboo structure formation. Scale bar is 500 nm in (a) and 200 nm in (b). Optimized set of triangles's extensions selected for further experiments is RV 12A.

Supplementary Figure 10: Data analysis of dried silica-coated DNA origami nanotube arrays on $Si/SiO_2$ on wafers

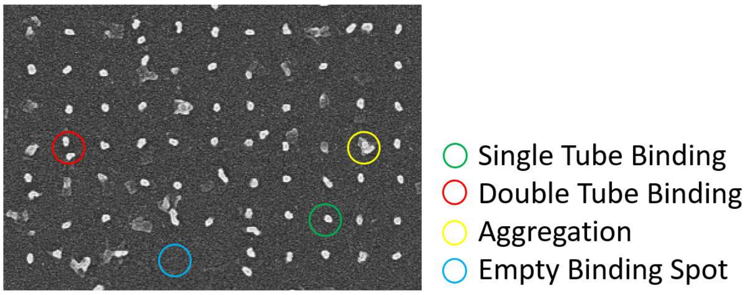


SEM image of $Si/SiO_2$ wafer with silica-coated nanotubes annealed to the triangles bearing 27 12A extensions arranged in lattice with period of 250 nm. Examples of binding events are marked: single standing tube (green), multiple tubes (red), tube/triangle aggregate (yellow), empty binding site (blue).

Supplementary Figure 11: Dried silica-coated nanotube arrays as a function of number and geometry of 12A extensions on triangles

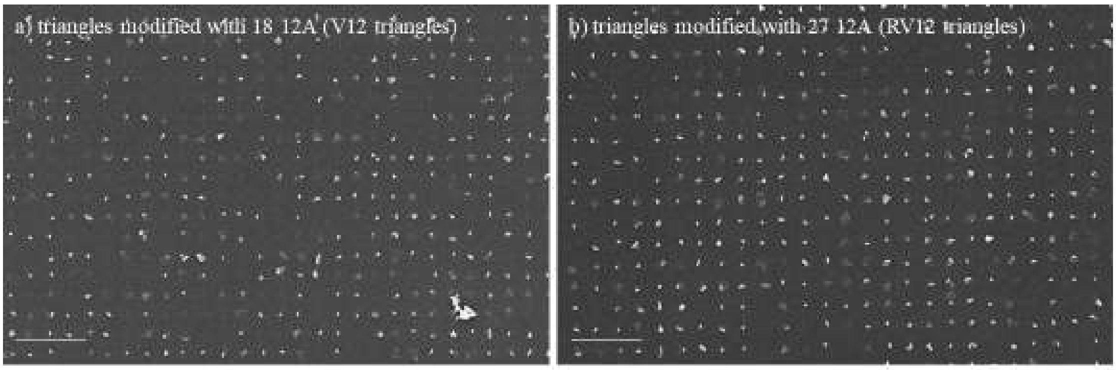


c)

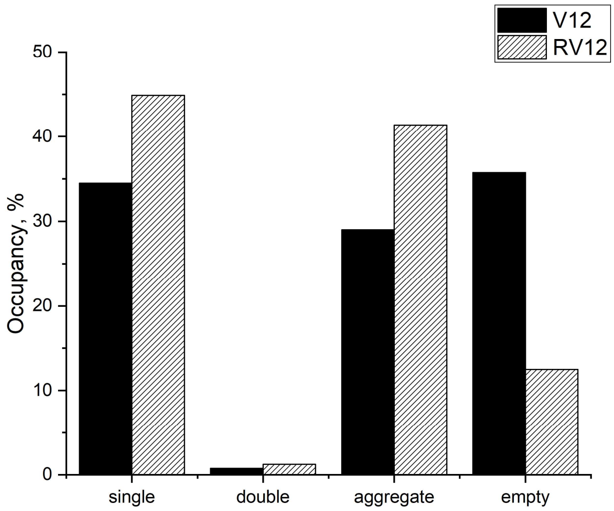


Annealing of tubes modified with 48 11T extensions (11T tubes) to triangles modified with 18 12A (V12 triangles) and 27 12A (RV12 triangles), arranged on the triangle as shown in Supplementary Figure 2. (a) SEM top-view image (scalebar 1 um) of silica-coated 11T tubes annealed to pre-patterned V12 triangles (b) SEM image top-view image (scalebar 300 um) of 11T tubes hybridized to patterned RV12 (c) Histograms with binding statistics for 11T tubes hybridized to V12 and RV12 triangles. 600 binding sites were scored for each sample. RV12 triangles were chosen for further experiments.

Supplementary Figure 12: Dried silica-coated nanotube arrays as a function of nanotube concentration

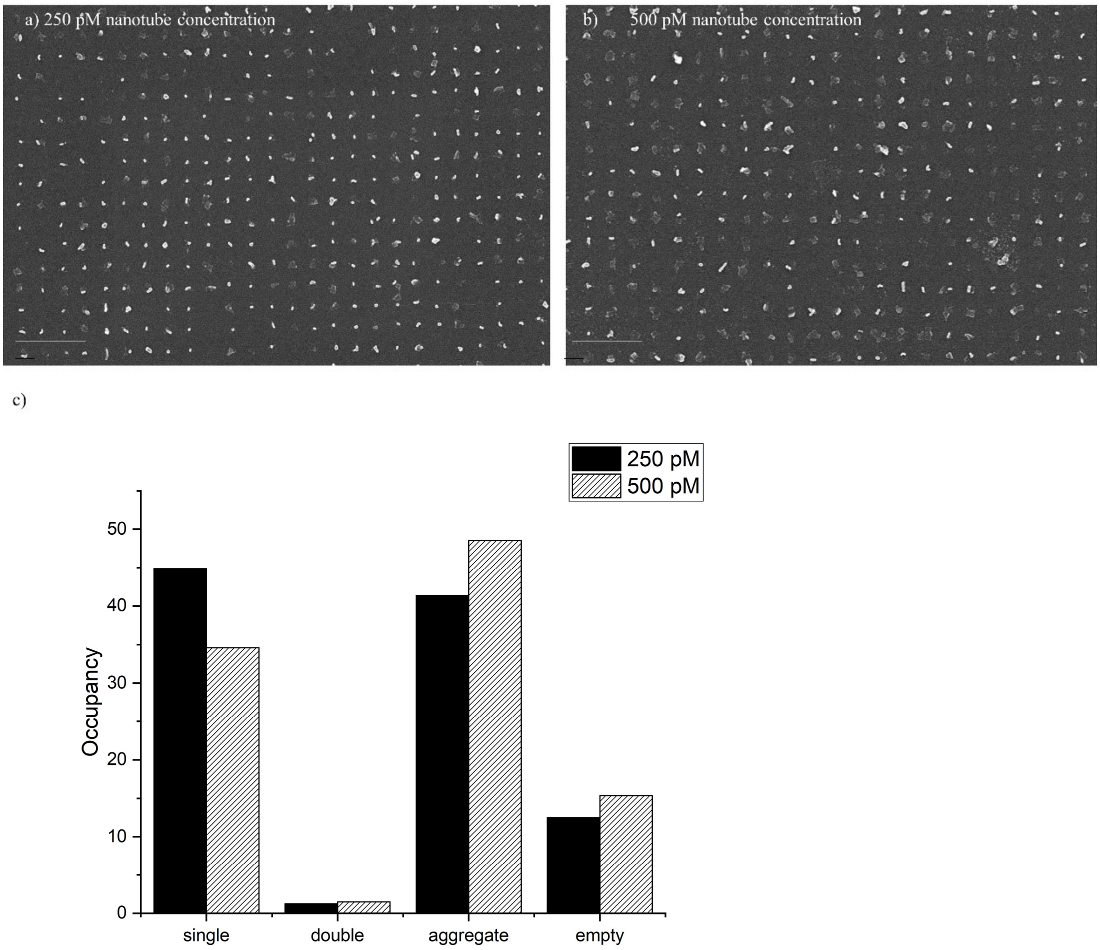


Annealing of tubes modified with 48 11T extensions (11T tubes) to triangles modified with 27 12A extensions as a function of nanotube concentration. SEM top-view image of silica-coated 11T tubes annealed to pre-patterned RV12 triangles at a concentration of 250 pM (a) or 500 pM (b) on $Si/SiO_2$ chips. Scalebar is 300 nm in (a) and 200 nm in (b). (c) Histograms with binding statistics for a 11T tube concentration of 250 pM and 500 pM. 600 binding sites were scored for each sample. Nanotube concertation of 250pM was chosen as optimal for further experiments.

Supplementary Figure 13: Dried silica-coated nanotube arrays as a function of $MgCl_2$ concentration in Hybridisation buffer

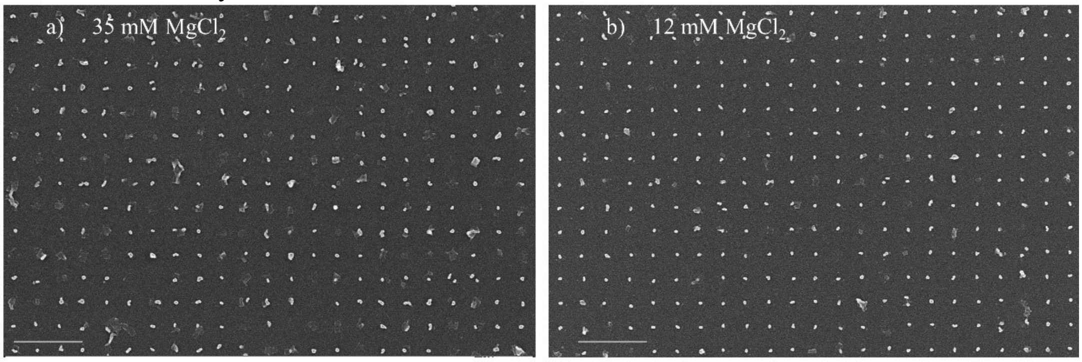


c)

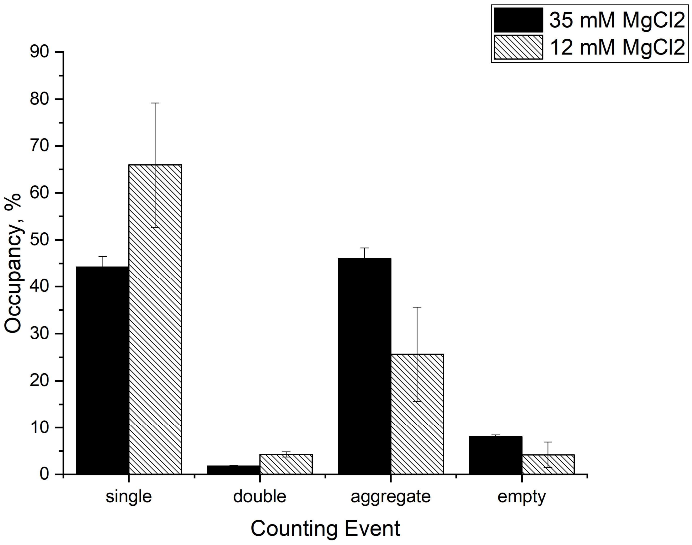


Annealing of tubes modified with 48 11T extensions (11T tubes) with concentration of 250 nM to triangles modified with 27 12A extensions (RV12 triangles) as a function of $MgCl_2$ concentration in Hybridisation buffer. (a) SEM top-view image (scalebar 1000 nm) of silica-coated 11T tubes annealed to pre-patterned 27 12A triangles at a concentration of 35 mM $MgCl_2$ or (b) 12 mM $MgCl_2$. (c) Histograms with binding statistics as a function of $MgCl_2$ concentration in Hybridisation buffer. Error bars are SD for N = 2 independent replications (of placement, washing, etc.) using different $Si/SiO_2$ chips from the same wafer. 600 binding sites were scored for each replicate.

Supplementary Figure 14: Hybrid silica-DNA nanotubes tubes in arrays on $Si/SiO_2$ wafer

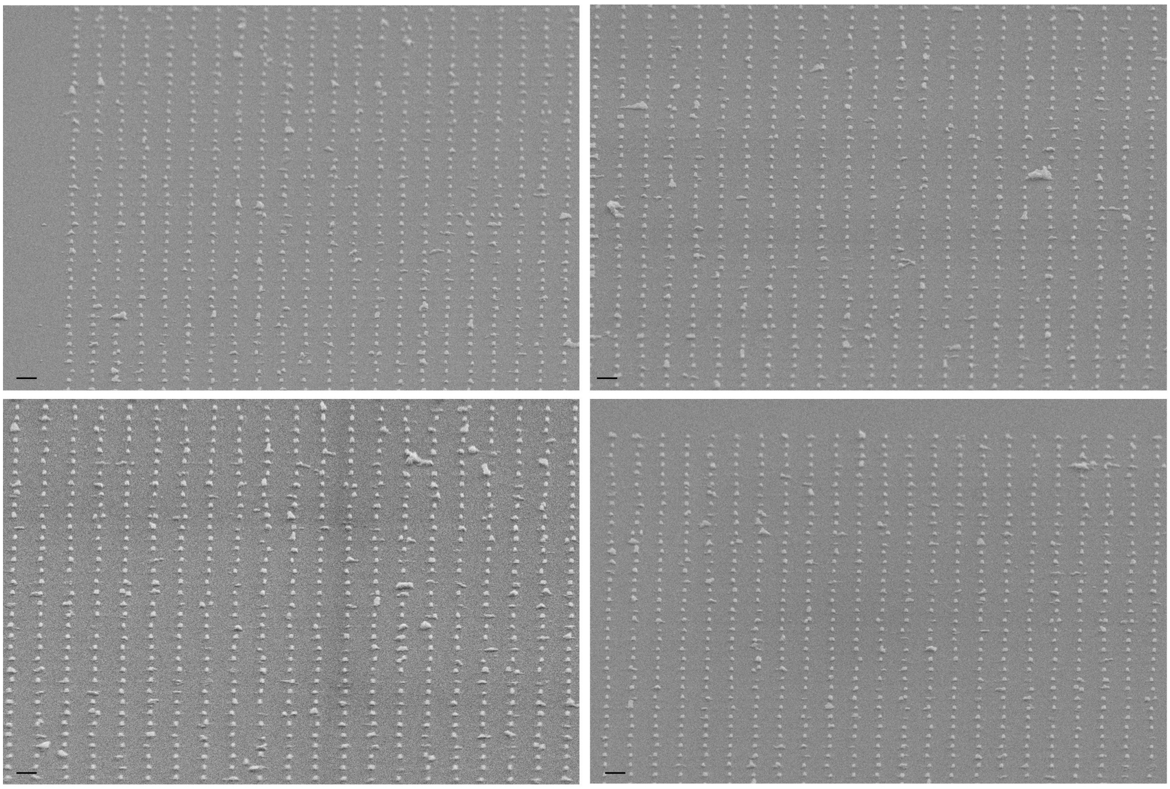

Tilted SEM image (scalebar 200 nm) of randomly selected areas of silica-coated 11T tubes annealed to pre-patterned 27 12A triangles at 12 mM $MgCl_2$ recorded on different spots of the same $Si/SiO_2$ chip.

Supplementary Figure 15: Wall thickness of hybrid silica DNA tubes in arrays

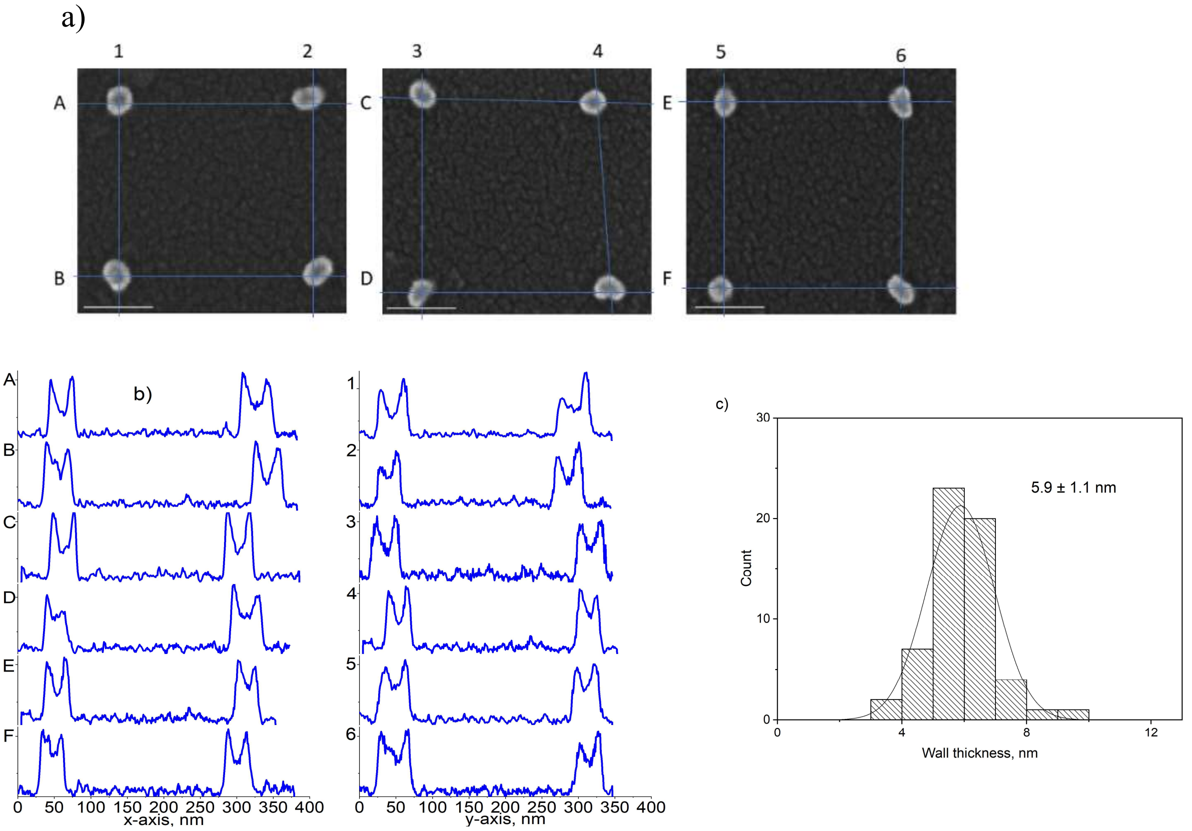


a) SEM top-view images (scalebar 100 nm) of silica-coated 11T tubes annealed to pre-patterned 27 12A triangles at 12 mM $MgCl_2$ with labelled scan lines. Scale bar is 100 nm. (b) The corresponding SEM line-scan profiles along the *x* and *y* axis (c) SEM measurement of the wall thickness.

Supplementary Figure 16: AFM images of Si/$SiO_2$ wafers patterned with triangles

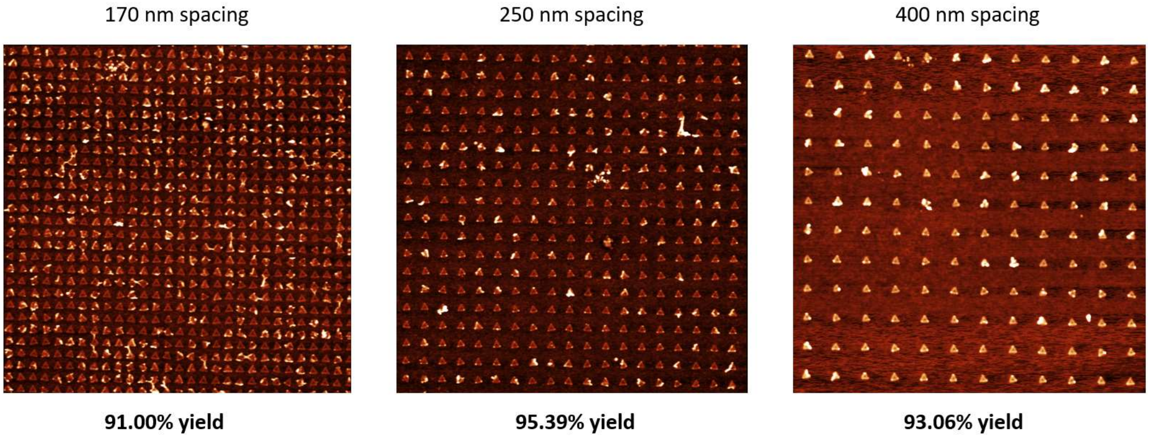


Dry-mode AFM images of Si/$SiO_2$ wafers patterned with triangles bearing 27 12A extensions. 300 events were counted for the 170 nm spacing, 304 events for the 250 nm spacing and 144 for the 400 nm spacing. Different periods of 170 nm, 250 nm, 400 nm between triangles result in comparable single triangle binding yields of 91%, 95%, 93%, respectively.

Supplementary Figure 17: Dried silica-coated nanotube arrays as a function of array period

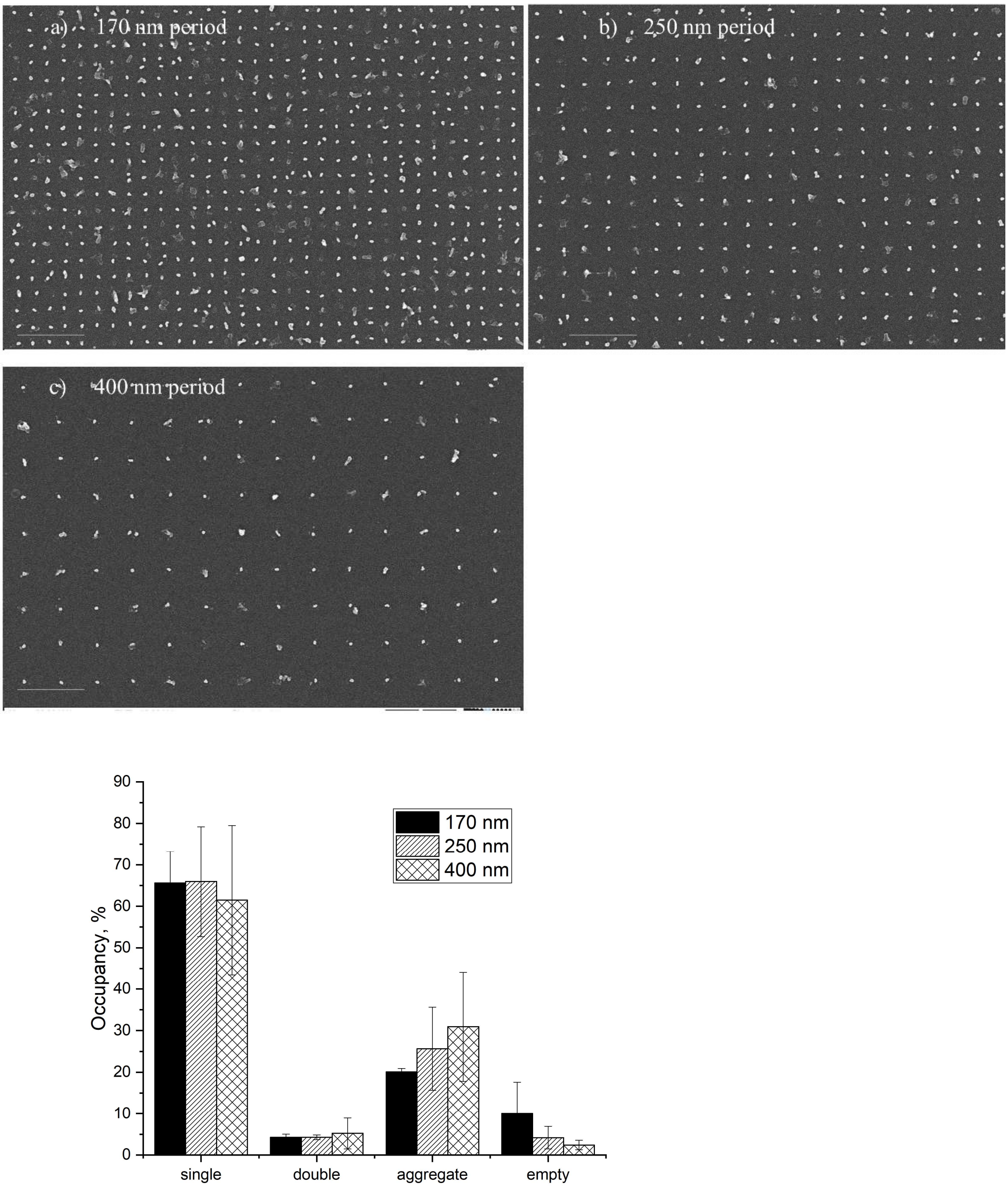


Annealing of tubes modified with 48 11T extensions (11T tubes) with a concentration of 250 nM to triangles modified with 27 12A extensions (RV12 triangles) as a function of array period (170 nm, 250 nm, 400 nm). SEM top-view image (scalebar 1000 nm) of silica-coated 11T tubes annealed to 27 12A triangles pre-patterned on Si/SiO2 chips in square array with 170 nm (a), 250 nm (b) and 400 nm (c) periods. (d) Histograms with binding statistics as a function of array period. Error bars are SD for N = 2 independent replications (of placement, washing, etc.) using different chips from the same wafer. 600 binding sites were scored for each replicate.

Supplementary Figure 18: AFM image of glass patterned with triangles

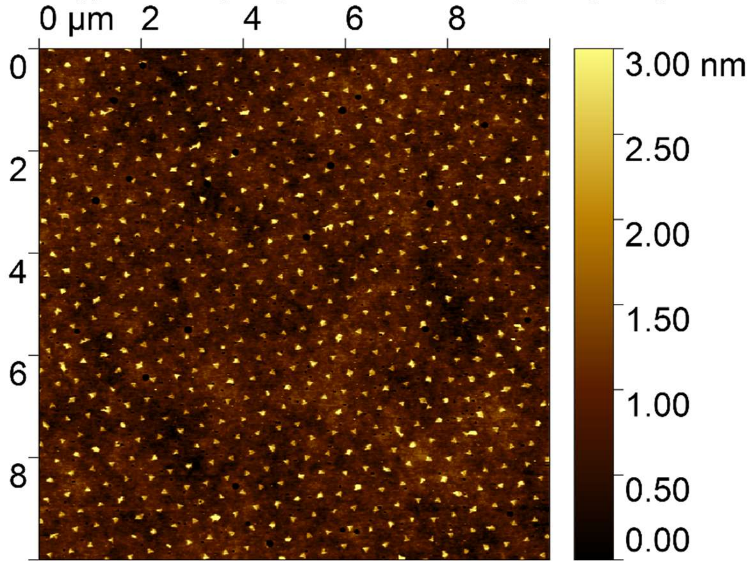


Dry-mode AFM images of glass patterned with triangles bearing 27 12A extensions. Ssingle triangle binding yield is 95%. 600 events were counted in a single experiment.

Supplementary Figure 19: SEM image of glass patterned with triangles and tubes as a function of nanotube incubation time

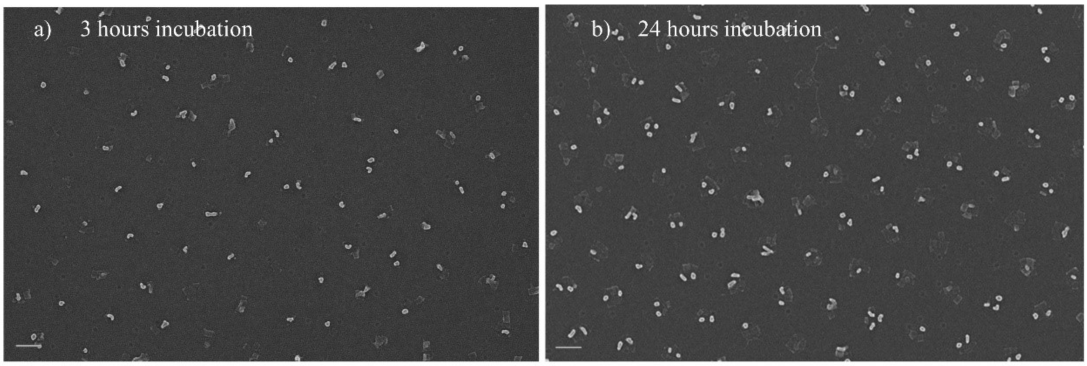


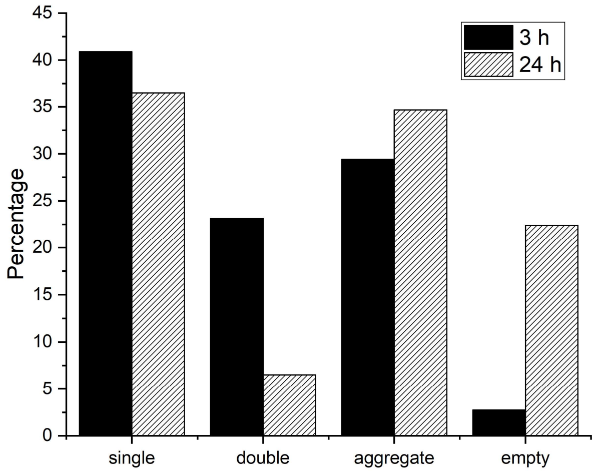


Annealing of tubes modified with 48 11T extensions (11T tubes) with concentration of 300 nm to triangles modified with 27 12A extensions (RV12 triangles) on glass. (a) SEM top-view image (scalebar 200 nm) of silica-coated 11T tubes annealed to 27 12A triangles pre-absorbed on glass using method of nanosphere lithography. Nanotube incubation time is 3 h (a) and 24 h (b). (c) Histograms with binding statistics as a function of function of incubation time. 600 binding sites were scored for each sample.

Supplementary Figure 20: SEM image of glass patterned with triangles and tubes

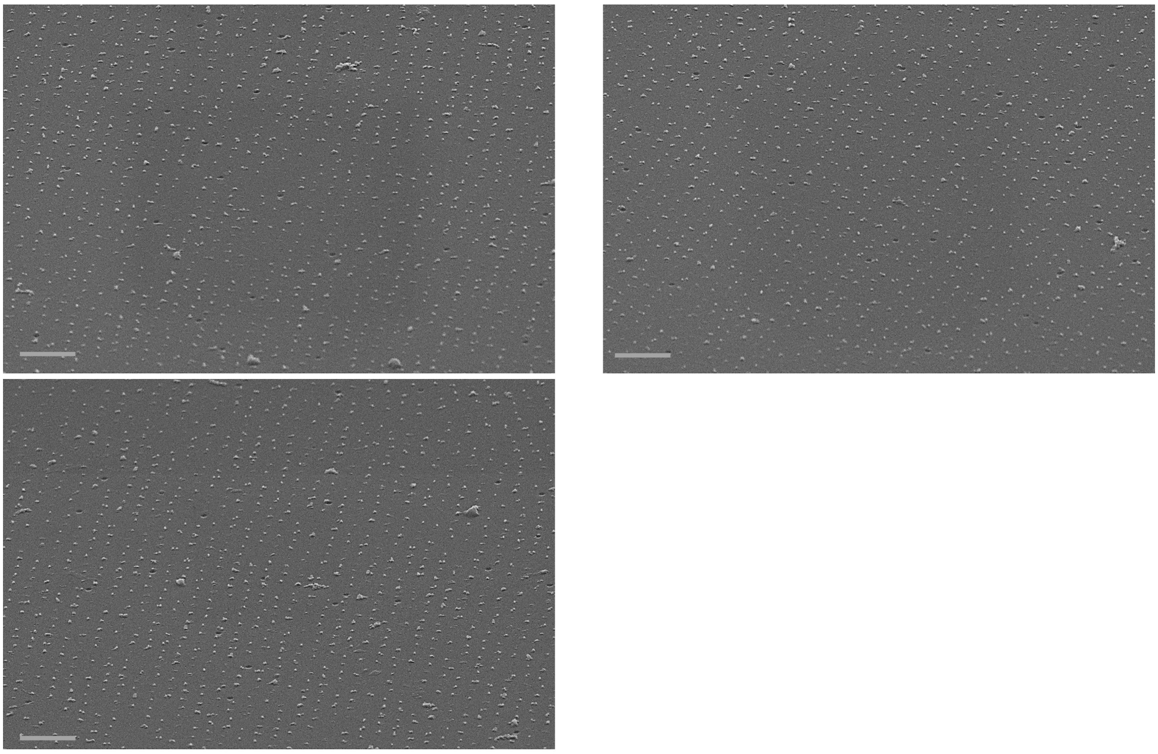

Tilted SEM image (scalebar 200 nm) of randomly selected areas of silica-coated 11T tubes annealed to pre-patterned 27 12A triangles at 12 mM $MgCl_2$ recorded on different spots of the same glass chip. Images are recorded at the middle point of the glass chip, and in points, 1 mm away from the middle point.

Supplementary Figure 21: TEM images of the barrels

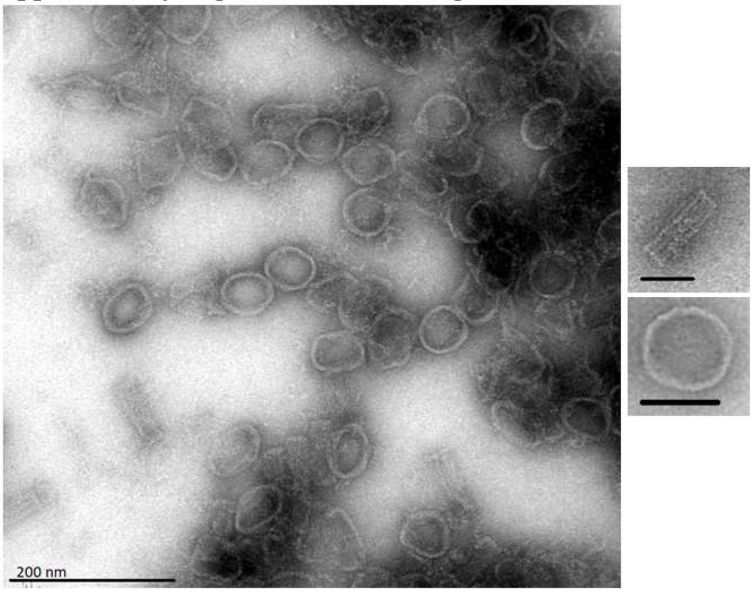


TEM images of DNA-origami barrels after ultracentrifugation and Amicon purification. Scale bars in zoomed-in images are 50 nm.

Supplementary Figure 22: Data analysis of dried silica-coated DNA origami barrel arrays on Si/$SiO_2$ wafers.

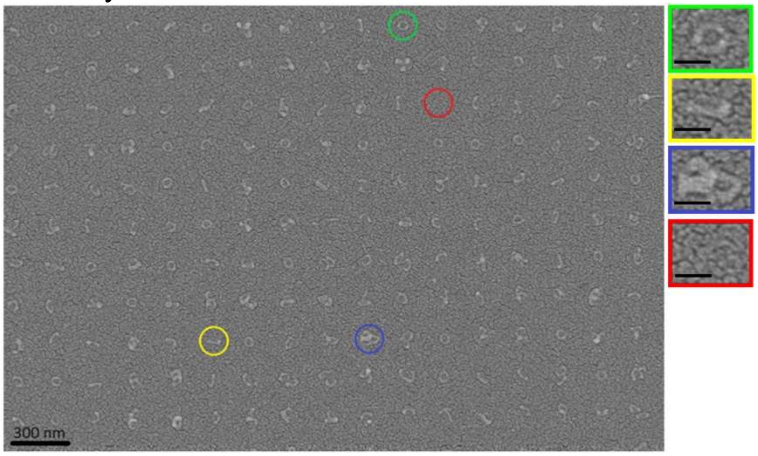


SEM images of Si/$SiO_2$ wafers with barrels arranged in lattice with period of 200 nm. Placement is performed on circular binding sites of 55 nm diameter, exposed at 400 uC/cm$^2$ dosage, 200 pM origami concentration. Examples of binding events are marked: single binding (green), single barrels lying sideways (yellow), multiple binding (blue), and empty binding sites (red). Scale bars in the zoomed-in images are 60 nm.

Supplementary Figure 23: DNA origami barrel arrays on $Si/SiO_2$ wafers as a function of barrel concentration.

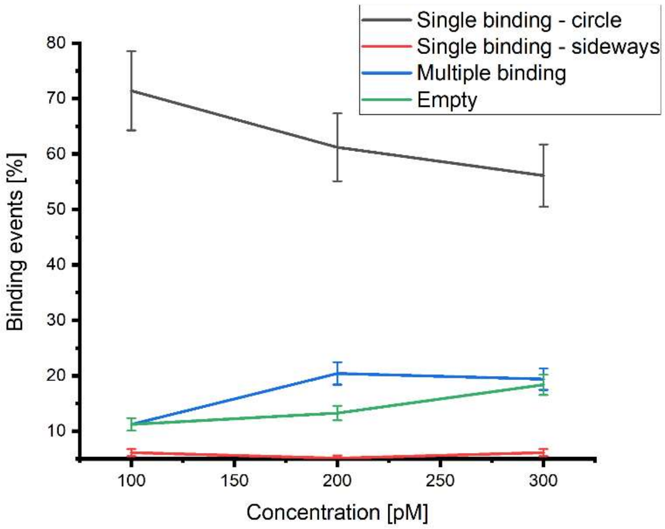


Variation of Concentration: Percentage of binding sites that show a certain event vs. the varied concentration. Lines are drawn to guide the eye. Placement on circular binding sites of 50 nm diameter, exposed at 400 $uC/cm^2$ dosage. 600 binding sites were scored for each sample.

Supplementary Figure 24: DNA origami barrel arrays on $Si/SiO_2$ wafers as a function of e-beam dosage

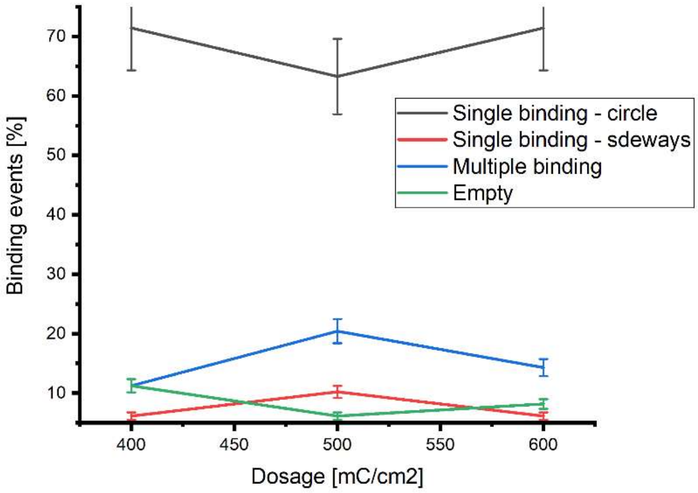


Variation of Dosage: Percentage of binding sites that show a certain event vs. the varied dosage. Lines are drawn to guide the eye. Placement of 100 pM barrels on circular binding sites of 50 nm diameter. 600 binding sites were scored for each sample.

Supplementary Figure 25: DNA origami barrel arrays on Si/$SiO_2$ wafers as a function of binding site size

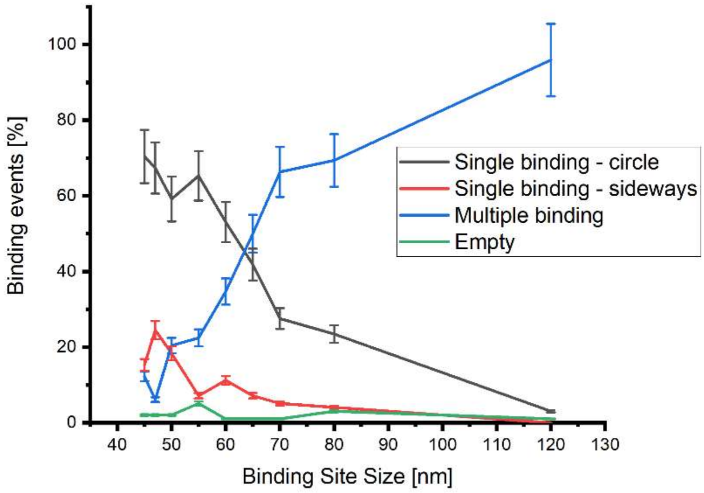


Percentage of binding sites that show a certain event as a function of the circular binding site diameter. Lines are drawn to guide the eye. Placement of 100 pM barrels on circular binding sites of diameters 45 nm to 60 nm, 300 pM on sites 65 nm to 120 nm, all exposed at 400 uC/cm$^2$. 600 binding sites were scored for each sample.

Supplementary Figure 26: Optimal barrel placement

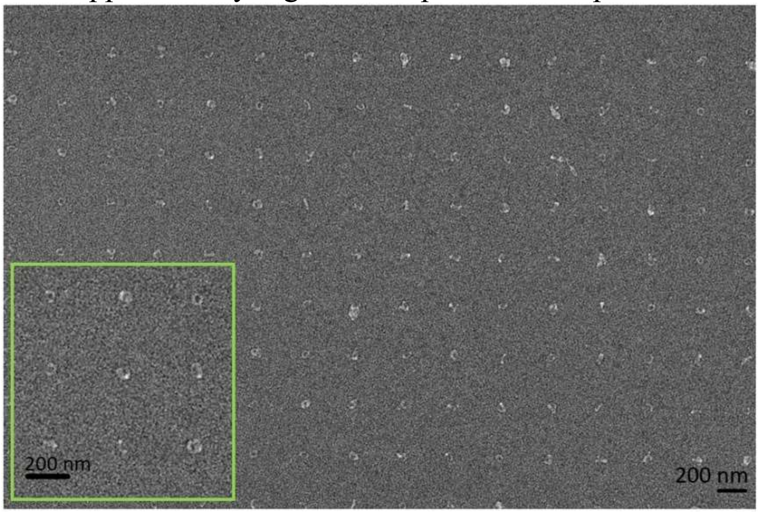


Optimal barrel placement. 100 pM barrels on circular binding sites of 45 nm diameter, exposed at 400 uC/cm$^2$.

Supplementary Figure 27: DNA Origami tetrapod design

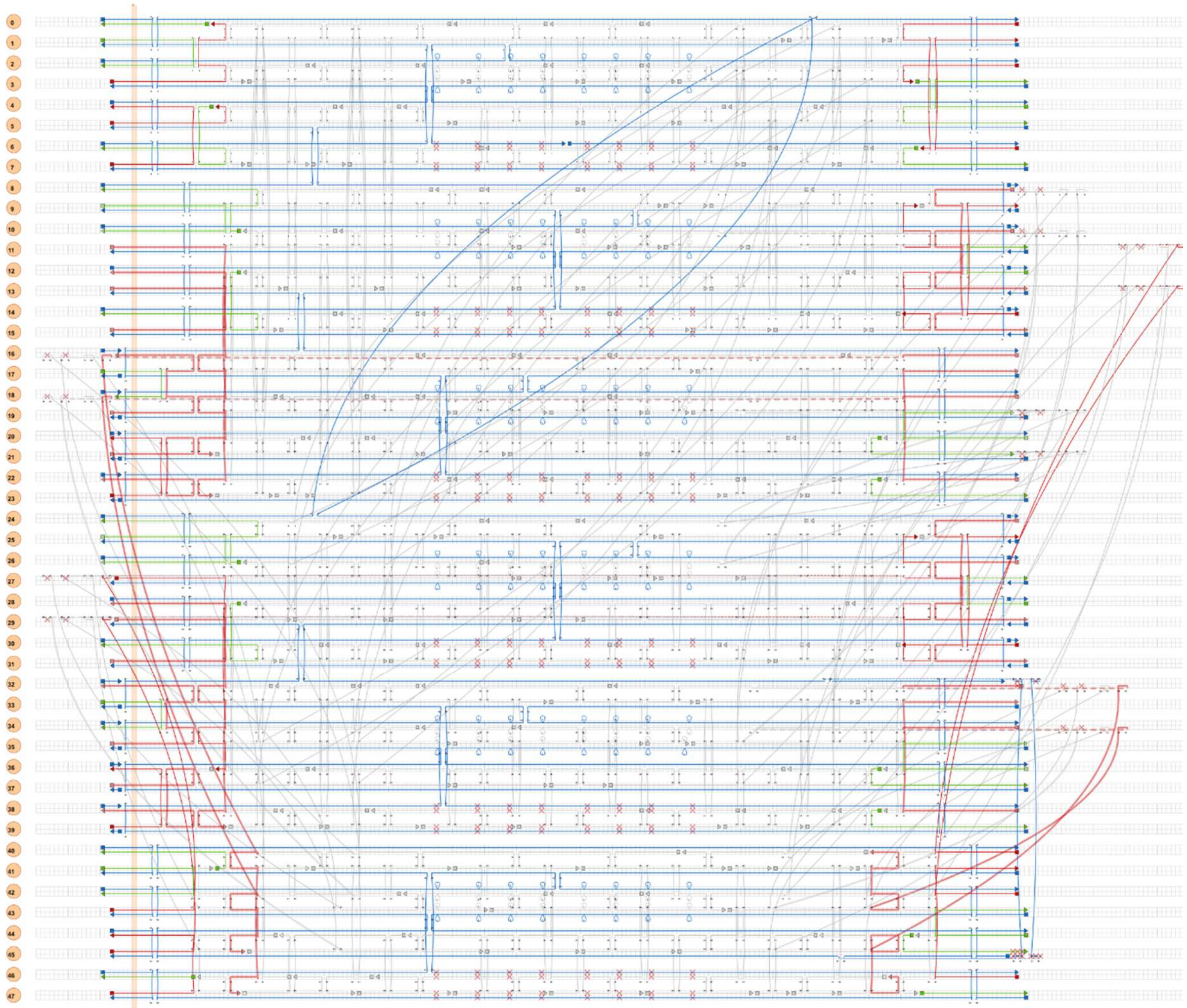

CaDNAno layout of the DNA origami tetrapod structure design.

Supplementary Figure 28: TEM image of the tetrapods

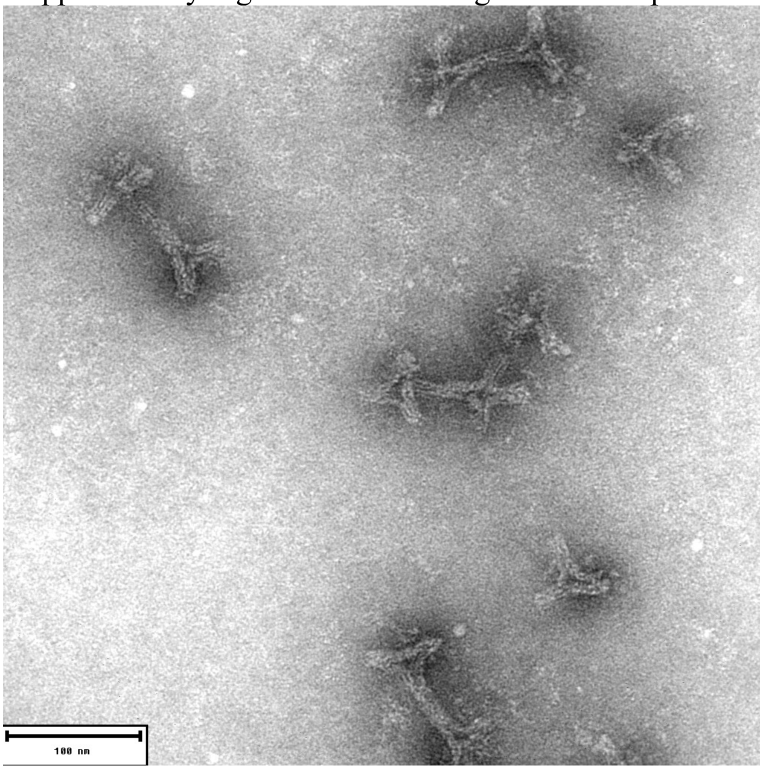


TEM images of DNA origami tetrapods after Gel Purification. Scalebar: 100 nm.

Supplementary Figure 29: Data analysis of dried silica-coated DNA origami tetrapod arrays on $Si/SiO_2$ wafers

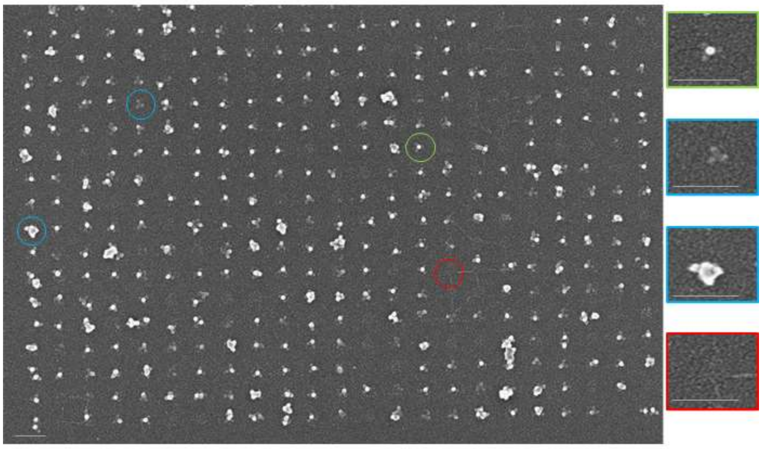

SEM images of $Si/SiO_2$ wafers with tetrapods arranged in lattice with period of 200 nm. Placement is performed on square binding sites of 35 nm diameter, exposed at 400 uC/cm$^2$ dosage, 400 pM origami concentration. Examples of binding events are marked: single binding "on three legs" (green), multiple binding/aggregated or deformed tetrapod (blue), and empty binding sites (red). Scale bars are 200 nm.

Supplementary Figure 30: DNA origami tetrapod arrays on Si/$SiO_2$ wafers as a function of square binding site size

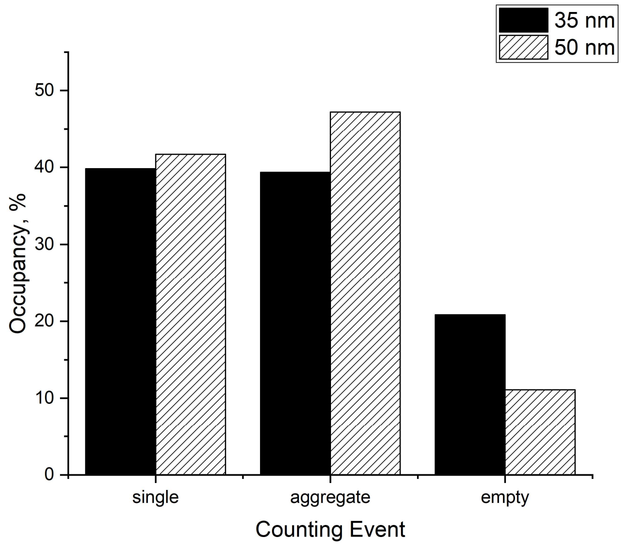


Percentage of binding sites that show a certain event as a function of the square binding site size. Placement of 400 pM barrels on binding sites with size of 35 nm and 50 nm, all exposed at 400 uC/$cm^2$. 600 binding sites were scored for each sample. Binding sites with the size of 35 nm were chosen for further experiments.

Supplementary Figure 31: DNA origami tetrapod arrays on Si/$SiO_2$ wafers as a function of e-beam dosage

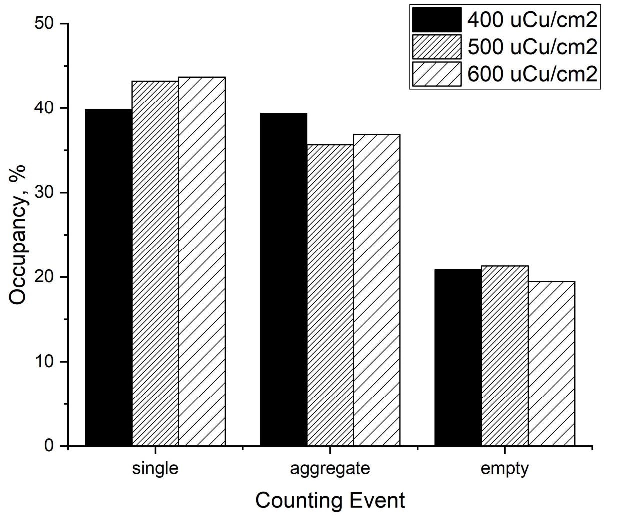


Variation of Dosage: Percentage of binding sites that show a certain event vs. the varied e-beam dosage. Placement of 400 pM tetrapods on square binding sites of 35 nm diameter. Dosage of 600 uCu/$cm^2$ was chosen for further experiments. 600 binding sites were scored for each sample.

Supplementary Figure 32: Optimal tetrapod placement

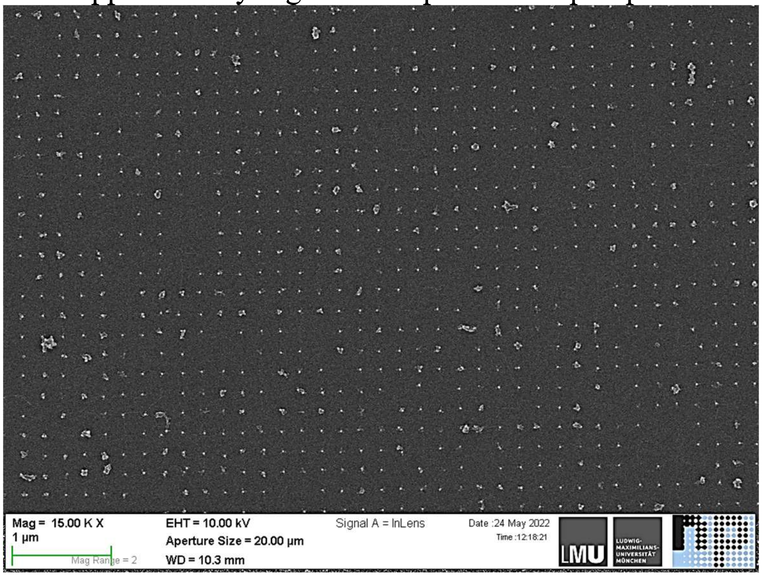


Optimal tetrapod placement. 400 pM tetrapods are deposited to square binding sites of 35 nm size, exposed at 400 uC/cm$^2$.

Supplementary Figure 33: DNA Origami 24HB design

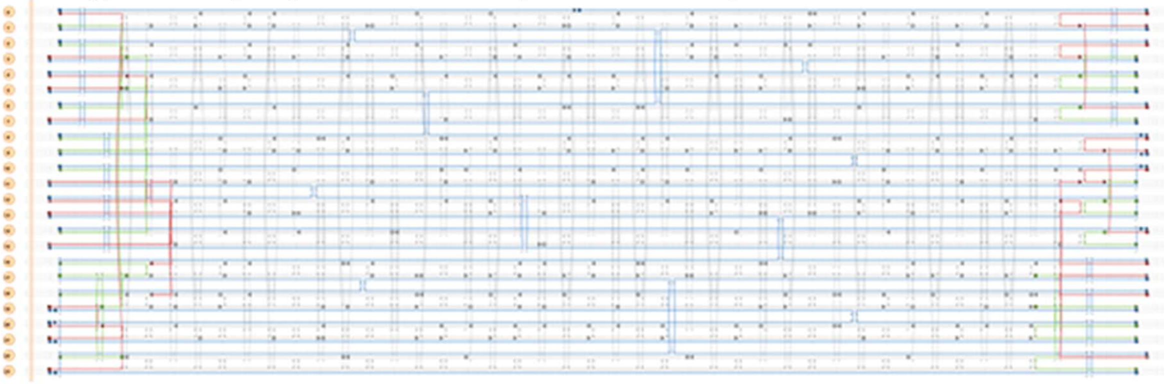

CaDNAno layout of the DNA origami 24HB structure design.

Supplementary Figure 34: Design of the interface between tetrapod and 24HB

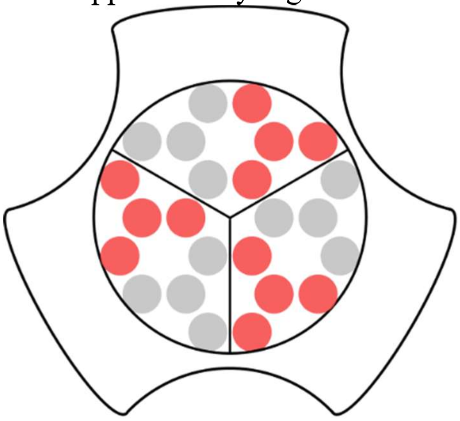

Shown is one of the end surfaces of the arms of the tetrapod with a 24HB cross-section. Out of the 24 available endings of helices on each surface, 12 are extended with single-stranded DNA linkers (marked as red circles). Hybridization between these tetrapod linkers and complementary anchor strands extending from the 24HB'ends connects end surfaces of the tetrapod "legs" and 24HBs.

Supplementary Table 1: Design Parameters of DNA Origami Nanotube

| Nanotube parameter | Description | Value |
|---|---|---|
| Number of duplexes $N_d$ | Number of double-stranded DNA duplexes aligned side-by-side and held together with crossovers. | 48 |
| Dihedral angle $\theta_{pleat}$ | The angle between duplex $n$ and its two neighbours relative to the "inside" surface of the curved sheet or tube. | 171.4$^{o}$ |
| Intrinsic curvature $\theta_{curve}$ | The angle formed between three even/odd duplexes relative to a flat sheet. | 17$^{o}$ |
| Diameter $D$ | $D = \frac{2\, ihDist \sin(\pi - \frac{\pi\, \theta_{pleat}}{360} - \frac{2\pi}{N_d})}{\sin(\frac{2\pi}{N_d})}$, where $\theta_{pleat}$ is the dihedral angle, $N_d$ is the number of duplexes and $ihDist$ is the interhelical distance in nanometres, 2.7 nm, a value measured from in-solution studies of one-layer DNA origami structure with square lattice type [7]. | 41.3 nm |
| Height $H$ | Nanotube height in base pair (bp) and nm, assuming 0.34 nm DNA helix rise per bp [8]. | 149 bp (~50 nm) |

Supplementary Table 2: Buffers

| | |
|---|---|
| Folding buffer | 10 mM Tris, 1 mM EDTA, 12.5 mM $MgCl_2$, pH 8.35 |
| $SiO_2$ Placement buffer | 5 mM Tris, 35 mM $MgCl_2$, pH 8.35 |
| $SiO_2$ Hybridisation buffer | 5 mM Tris, 12.5 mM $MgCl_2$, pH 8.35 |
| $SiO_2$ Imaging buffer | 5 mM Tris, 35 mM $MgCl_2$, pH 9 |
| Glass Placement buffer | 40 mM Tris, 40 mM $MgCl_2$, pH 8.35 |
| Glass Imaging buffer | 10 mM Tris, 35 mM $MgCl_2$, pH 8.9 |
| Silica coating buffer | 40 mM Tris, 2 mM EDTA-2Na, 12.5 mM $MgAc_2$, pH 8.0 |
| Silica coating buffer with 4% pre-hydrolyzed silica precursors | Silica coating buffer with 2% -trimethoxysilylpropyl-N,N,N-trimethylammonium (TMAPS) (50% (wt/wt) in methanol) and 2% tetraethylorthosilicat (TEOS). |

### Materials and Methods

#### DNA Origami design, preparation and purification

DNA origami nanotubes and tetrapods were designed using the software caDNAno [2]. Details of the design of DNA nanotubes can be found in Supplementary Note 1. Details of the design of DNA tetrapods and 24HBs can be found in Supplementary Note 7 and Supplementary Note 9, respectively.

Design-specific staple strands were purchased from IDT Technologies, the scaffold strands (p7249, p8634) were produced from M13 phage replication in Escherichia coli. All chemicals were obtained from Sigma Aldrich, unless otherwise stated. DNA origami used in this work (nanotubes, triangles, barrels, tetrapods and 24HB) were folded by mixing scaffold strands with excess of staple starnds (and miniscaf short scaffold-parity strands in case of DNA origami barrels) in folding buffer (buffers used in the work can be found in Supplementary Table 1). Samples were thermally annealed in a PCR machine (Tetrad 2 Peltier thermal cycler, Bio-Rad) and purified from excess staples by methods of Amicon filtration [9] (triangles, nanotubes), or PEG precipitation [10] (tetrapods, 24HB) or ultracentrifugation [11] (barrels). Full description of folding and purification of each type of DNA origami can be found below.

#### Folding and purification of DNA Origami nanotubes

Scaffold DNA was mixed with all staples except for 48 edge staples that were replaced by 3′ 11(T) modified staples (HPLC purified, Biomers) in Folding buffer (10 mM Tris pH 8.4, 1 mM EDTA, 12.5 mM $MgCl_2$, see Supplementary Table 1, at the following final concentrations:

| Component | Target Concentration (nM) | Excess |
|---|---|---|
| Scaffold 7249 | 20 | 1 |
| Staples core mix without 48 edge staples | 100 | 5 |
| 48 edge staples polyT-modified | 400 | 20 |

The folding mixtures (50 µL) were subjected to the following nonlinear thermal annealing ramp in a PCR machine:

| Temperature (°C) | Time per °C (min) |
|---|---|
| 65 | 15 |
| 60-20 | 3 min 12 sec |
| 4 | storage |
| Total duration: ~3 h | |

After folding, nanotubes were purified from excess staples using 100 kD molecular weight cut-off filters (Amicon Ultra-0.5 Centrifugal Filter Units with Ultracel-100 membranes) [9]. We followed to protocol below:

- Wet the filter by adding 500 µL of Folding buffer.
- Spin filter at 8000 rcf for 5 min at RT down to 30 µL. Discard the filtrate.
- Add 200 µL of unpurified origami and 250 µL of Folding buffer. Spin at 8000 rcf for 7 min at RT.

- Discard the filtrate. Add 450 µL Folding buffer and spin at 8000 rcf for 7 min at RT.
- Repeat previous step three more times.
- Invert the filter onto a clean tube and spin at 2000 rcf for 3 min to collect purified origami (~ 30µL).

After purification, UV–vis absorption spectra of samples were recorded with NanoDrop spectrophotometer (Thermo Scientific) in order to determine DNA origami concentration using the molar extinction coefficient of the DNA origami as that of a p7249 dsDNA molecule.

The concentration of nanotubes after Amicon purification is usually 100 nM, recovery is 70-80%. Stock solution of nanotubes can be stored at 4ºC in LowBind DNA Eppendorf tubes.

**Folding and purification of DNA Origami triangles**

Scaffold DNA (p7249) was mixed with all staples except for 27 modified staples that were replaced by 5′ (T) modified staples (HPLC purified, Biomers) in folding buffer (10 mM Tris pH 8.4, 1 mM EDTA, 12.5mM $MgCl_2$) at the following final concentrations:

| Component | Target Concentration (nM) | Excess |
|---|---|---|
| Scaffold 7249 | 20 | 1 |
| Core Mix | 100 | 5 |
| Up to 27 staples polyA-modified | 400 | 20 |

The folding mixtures (50 µL) were subjected to the following nonlinear thermal annealing ramp:

| Temperature (°C) | Time per °C (min) |
|---|---|
| 65 | 15 |
| 64-60 | 5 |
| 59-40 | 45 |
| 39-36 | 30 |
| 35-20 | 5 |
| 4 | storage |
| Total duration: ~18 h | |

Folded samples were purified to remove excess staple strands via Amicon filtration as described above for the nanotubes. The concentration of triangles after Amicon purification is usually 70 nM, recovery is 40-50%.

**Folding and purification of DNA Origami barrels**

The design of the barrels with a diameter of 60 nm and a height of 27 nm is taken from [12]. The structure is designed in a honeycomb lattice and is based on a 2D rectangular DNA sheet with horizontally arranged helices.

Folding of DNA origami was performed as described before in [12] for 60-27 nm barrel monomers. Briefly, scaffold DNA (p7249) was mixed with staples strands, mini-scaffold strands and coaxial top and bottom staples in Folding buffer with 10 mM $MgCl_2$.

| Component | Target Concentration (nM) | Excess |
|---|---|---|
| Scaffold p7249 | 20 | 1 |
| Core Mix (core staples and miniscaffold staples) | 200 | 10 |
| Coaxial staples top and bottom | 200 | 10 |

The folding mixtures (50 µL) were subjected to the following nonlinear thermal annealing ramp:

| Temperature (°C) | Time per °C (min) |
|---|---|
| 65 | 15 |
| 50-40 | 10 h |
| 4 | storage |
| Total duration: ~66 h | |

After the folding, the DNA-origami barrels were purified from excess staples in two separate steps. First, a rate-zonal centrifugation procedure [11] is performed using a Beckman Coulter Optima MAX-XP Ultracentrifuge. 30-60% glycerol gradients in Folding buffer with $MgCl_2$ concentration 12.5 mM are prepared by pipetting 80 µl of each glycerol concentration into an ultracentrifuge tube, starting with the highest glycerol concentration on the bottom of the tube. 160 µl of folded barrels are added on top. The tubes are then placed in the Beckman SW-Ti rotor and spun at 4°C at 300 000 rcf for 15 min. Following the ultracentrifugation, the sample is divided into nine fractions of 80 µl. They are carefully taken from the top and named fractions 1-9 in the order in which they are picked up and loaded to a 2% Agarose Gel (data not shown). Fractions 5-9 are collected for further experiments, and purified from glycerol using a method of Amicon centrifugation following the protocol:

- Wet the filter by adding 500 µL of Folding buffer.
- Spin filter at 8000 rcf for 5 min at RT down to 30 µL. Discard the filtrate.
- Add 200 µL of barrels in Folding buffer/glycerol and 250 µL of Folding buffer. Spin at 8000 rcf for 7 min at RT.
- Discard the filtrate. Add 450 µL Folding buffer and spin at 8000 rcf for 7 min at RT.
- Invert the filter onto a clean tube and spin at 2000 rcf for 3 min to collect purified barrels (~ 30µL).

The concentration of barrels after purification is usually 10 nM and staples are no longer visible by agarose gel (Materials and Methods Figure 1).

Materials and Methods Figure 1. Gel electrophoresis image of barrels

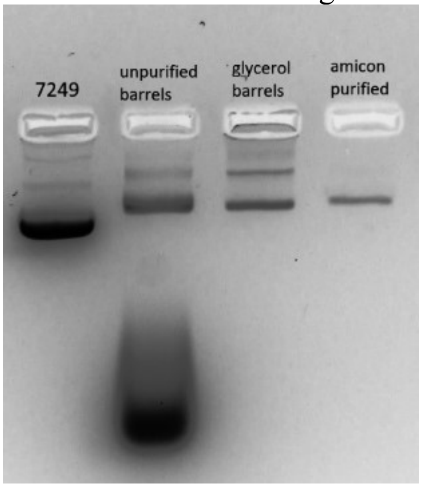


Gel electrophoresis image of barrels at different stages of the purification process: unpurified barrels after folding, barrels, purified by rate-zonal centrifugation (glycerol barrels), and barrels, purified by rate-zonal centrifugation and following Amicon filtration (amicon purified).

**Folding and purification of DNA Origami tetrapods**

The tetrapod structure is folded from p8634 scaffold (15 nM concentration) and around 200 staples (120 nM; ordered from IDT) in 10 mM Tris buffer with 1 mM EDTA (pH = 8.0; 1x TE buffer) and 20 mM $MgCl_2$ in 100 µL aliquots.

The folding protocol for the structure was optimized with light scattering-based monitoring of the folding [13]:

| Temperature (°C) | Time per °C (min) |
|---|---|
| 67 | 15 |
| 60-50 | 40 |
| 4 | storage |
| Total duration: 7 h 40 min | |

The folded structures were diluted to 1 mL volume by folding buffer (1x TE with 20 mM $MgCl_2$) and mixed with 1 mL of 15% PEG-800 with 500 mM NaCl and 1x TE buffer. The mixture was centrifuged for 30 min at 20000 RCF and 4°C. After centrifugation the supernatant was removed and the pellet resuspended in 1x TE with 5 mM $MgCl_2$.

**Folding and purification of DNA Origami 24HB**

The 24HB structure is folded from p7249 scaffold (15 nM concentration) and around 200 staples (120 nM; ordered from IDT) in 10 mM Tris buffer with 1 mM EDTA (pH = 8.0; 1x TE buffer) and 18 mM $MgCl_2$ in 100 µL aliquots.

The folding mixtures were subjected to the following nonlinear thermal annealing ramp in a PCR machine:

| Temperature (°C) | Time per °C (min) |
|---|---|
| 65 | 15 |
| 60-20 | 3 min 12 sec |
| 4 | storage |
| Total duration: ~3 h | |

After folding the structure was purified with PEG precipitation as described above for the tetrapods.

**Annealing of DNA origami in buffer solution**

For the triangular origami, 9, or 18, or 27 staples close to the middle hole were modified by extending 8, 12 or 20 adenine nucleotides on the 5′ end (Supplementary Figure 2). These $A_8$, $A_{12}$, $A_{20}$-modified DNA staples were introduced into the DNA scaffolds in place of the original DNA staples. After thermal cycling, excess staples were removed by filtering the origami solution through a 0.5 mL Amicon 100 kDa in a same way as described above.

For nanotube DNA origami, 24 staples on each edge of the tube were modified by extending 11 thymine nucleotides on the 3′ end (see Supplementary Note 2 and Supplementary Figure 2). These $T_{11}$-modified DNA staples were introduced into the DNA scaffolds in place of the original DNA staples and folded and purified as described above.

The effectiveness of binding of T-modified DNA nanotubes to A-modified DNA triangles in suspension was studied at a final concentration of tubes and triangles of ~5 nM and varying length and number of linkers. The solutions were mixed in Placement buffer (Table S1) in 50 µl aliquots, annealed 24 h either at 37 °C in the thermal cycler or at room temperature, and imaged by gel electrophoresis and TEM.

**Preparation of the substrate**

**Nanopatterning of $Si/SiO_2$ wafers with e-line lithography**

Patterned $Si/SiO_2$ substrates were prepared by adaptation of procedure from [6] with slight modification. Clean $Si/SiO_2$ wafers with 100 nm thermal oxide were primed with hexamethyldisilazane (HMDS) in a 4 L desiccator always keeping the surface contact angle after HMDS deposition of 70-75°. Binding sites were patterned into poly(methyl methacrylate) resist by electron-beam lithography and the developed areas cleaned with $O_2$ plasma for 10 s in a plasma cleaner (PICO). The resist was stripped by ultrasonication in *N*-methylpyrrolidone (NMP) at 50 C for 30 min. The substrates were briefly rinsed with 2-propanol, then dried in a nitrogen stream and used immediately. All steps were carried out in LMU cleanroom.

Since the mechanism of origami-site binding is very complex and small details of substrate fabrication can have a large effect on placement, we place step-by-step protocol of preparation of Si/SiO2 wafers with reproducible contact angle of 70-75°.

***1. Substrate preparation***

Although the original protocol is started with thermal growth of $SiO_2$ layer on Si wafer, we found that commercially available $Si/SiO_2$ wafers with 100 nm $SiO_2$ (Prime Si + dry $SiO_2$ wafer 4 inch, thickness = 525 ± 25 µm, (100), 1-side polished, p-type (Boron), 1 - 10 Ohm cm) from Microchemicals provide sufficient surface quality for the reproducible placement.

Prior the experiments, 4-inch wafer was diced to 1 cm x 1 cm chips. Then, the protective layer on the wafers is removed by washing them in Acetone and isopropanol (IPA) for 70 s and drying in a stream of $N_2$.

The chips are then put to a PICO plasma cleaner at an oxygen flow rate of 45 sccm and a power setting of 80 W for a duration of 5 min in order to generate surface silanols. Next, the surface of the wafers is dehydrated on a heating plate at 150°C for 5 min and placed in a 4 L desiccator containing HMDS (hexamethyldisilazane) vapor. HMDS priming on a $SiO_2$ layer will deposit a monolayer of the priming agent with carbon atoms (methyl groups) pointing at the surface. To achieve reproducible surface properties of the HMDS-primed wafers, we optimized the procedure of HMDS priming.

Immediately after dehydration on a hot plate, the wafers are put into a 4 L glass desiccator alongside a petri dish containing 10 ml of HMDS.

Many experimental parameters may significantly influence the quality of priming, as volume of desiccator, humidity and temperature in the cleanroom and freshness of used HMDS, as well as time before dehydration on a hot plate and HMDS priming. Depending on the freshness of the HMDS, the chips are kept in the desiccator for 3 – 40 minutes, before getting baked on a hotplate at 150°C for 30 minutes. Afterwards, the contact angles of a chips are measured with Krüss Advance in Sessile drop mode and manual adjustment of the base line. Contact angles lie in the range of 70°-75°.

Then wafers are covered with a resist, PMMA (PMMA.A2 EM Resist LTD). This is done by placing ten drops of PMMA on the chips and spin-coating it for 1 s at 800 rpm and for 70 s at 2500 rpm. This results in a resist layer with a thickness of approximately 100-120 nm. Then the bottom of the chips is cleaned using Acetone, and chips are baked at 180°C for 30 s on a hotplate.

#### 2. *E-Beam Lithography*

In this step, the binding sites in the resist are defined by e-beam lithography with a E-line SEM with a 20 keV beam and a dosage of 300-400-500 uC/cm$^2$ at 60-75 pA current.

An example of a binding sites design in shown in Materials and Methods Figure 2. It includes two crosses that are used as a fiducial markers for finding the binding sites on the chip and nanopatterned area of approximately 30 μm x 30 μm size.

#### 3. *Activation of Binding Sites*

In order to activate the binding-sites before DNA origami placement, the PMMA has to be stripped from the binding sites so that the hydrophilic silanol groups can be exposed. Therefore, the wafers are put to PMMA developer MIBK:IPA, (1:3) and the shorter fractions of PMMA from the exposed areas are washed out. Afterwards, the wafers are washed in isopropanol to stop the reaction and dried using a nitrogen gun. In the next step, the silanol groups at the binding sites are generated by $O_2$ plasma in a PICO plasma cleaner at an oxygen flow of 45 sccm and a power setting of 80 W for 10 s. The plasma etches through the trimethylsilyl layer created during the deposition of HMDS and exposes the silanol groups. The rest of the resist on the background is removed by incubating the wafers in n-methyl pyrrolidone (NMP) in a sonicator at 50°C for 30 min. Afterwards, the NMP is washed off the wafers with isopropanol to stop the reaction and dried with a nitrogen gun. As a result, the binding sites on the wafers are modified with hydrophilic silanol groups and the background with hydrophobic trimethylsilyl groups.

Materials and Methods Figure 2: Nanopattern design

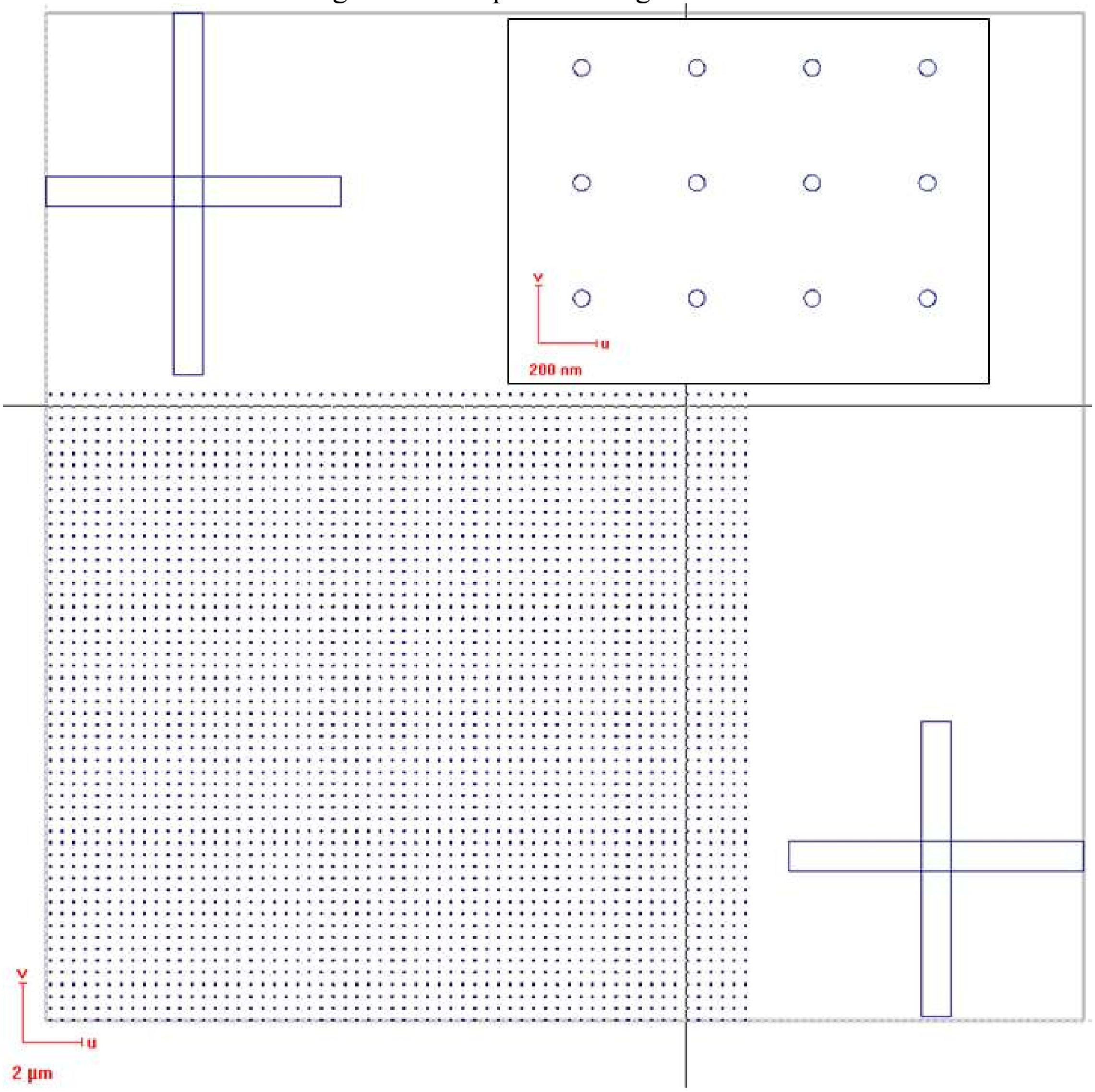


Schematic of a placement chip. Crosses are used as fiducial markers. (a) Nanopatterned area is usually up to 30 µm x 30 µm and consist of arrays of binding sites with different periods. Here, circular binding sites with diameter of 45 nm are arranged to the square lattice with 400 nm period. Insert shows the zoomed-in area of the lattice. (b) Area nanopatterned with fractal arrangement of 35-nm square binding sites, forming hexagonal lattice with side length of each hexagon of 170 nm. Insert shows the zoomed-in area of the pattern.

**Nanopatterning of glass wafers with nanosphere lithography**

Patterned glass substrates were prepared by adaptation of procedure from [14] without modification. A clean 1 $cm^2$ glass chips were purchased from Epredia. The PS nanospheres with diameter of 400 nm (Thermo Scientific™ Nanosphere™ Size standards 340A) were deposited by drop-casting onto an $O_2$-plasma activated glass chip surface and dried at a ~45° angle at R.T, forming a of a close-packed monolayer/multilayer of nanospheres. Then glass chips were primed with HMDS in a 1 L vapour chamber under vacuum. Then the PS nanosheres were lift off the surface by ultrasonication in water at RT for 1 min. Finally, the surface was blown dry with a nitrogen "gun" and baked at 120 °C for 5 min to stabilize the HMDS on the surface, and used immediately.

**Placement of DNA origami**

Before every placement experiment in this work, we folded and purified fresh DNA origami stock solutions as described above. Purified DNA origami samples were usually stored in the LowBind Eppendorf tubes no longer than for a week prior the experiment.

All buffers used in the placement of DNA origami are prepared fresh shortly before the experiments.

Nanopatterned substrates are used for the placement not later than 12 h after activation.

Full step-by-step protocol of the placement of DNA origami triangles, nanotubes, barrels and tetrapods can be found below.

**Placement of DNA origami triangles on Si/$SiO_2$ chips**

The original placement protocol was taken from [6] and used without modifications:

- **Substrate preparation.** For the placement of DNA origami triangles, 120 nm same side triangular binding sites arranged in square lattice with 170 nm, 250 nm or 400 nm period, defined by e-beam lithography on a Si/$SiO_2$ wafers, as described in above.
- **Preparation of DNA origami.** Triangles modified with PolyA extensions are folded and purified as described in Materials and Methods Section "Folding of DNA origami".
- **Placement**. A 50 mm petri dish with a lid is prepared with a moistened kimwipe to limit evaporation. A piece of parafilm is stuck to the ground of the petri dish to avoid shifting of the wafer. DNA Origami triangles are diluted to the 300 pM concentration in Placement Buffer in the LowBind DNA Eppendorf tube. A drop of 20 uL of the solution is placed in the middle of the 1 cm x 1 cm chip. The chip is placed in the closed, humid petri dish and the origami solution is allowed to incubate on the wafer for 1 h. After incubation, excess origami is washed away with at least 8 Placement buffer washes by pipetting 60 uL of fresh Placement buffer onto the wafer, and pipetting 60 uL off of the wafer. Each of the 8 washes consists of pipetting the 60 uL volume up and down 3-4 times to mix the fresh buffer with the existing buffer on the wafer. Then, the wafer is washed **5 times** using a buffer with **0.5%** Tween 20. After the 5th wash, the wafer is left to incubate for 30 minutes. Lastly, the wafer is buffer-washed with imaging buffer until all Tween is washed away and the drop has the original size and shape again. Prior drying, the wafer is incubated in the Imaging buffer for 2 h.
- **Drying.** The wafer is dried with an Ethanol gradient drying series: It is dipped in petri dishes filled with Ethanol mixtures (25%, 50%, 70%, 80%, 90%) for 10 seconds each and then left to air-dry. After drying, AFM images are taken in dry tapping mode with a Dimension Icon AFM (Bruker).

**Surface annealing of DNA origami nanotubes with triangles on Si/$SiO_2$ wafers**

- **Preparation of DNA origami.** We folded and purified fresh nanotubes modified with the polyT extensions and the triangles modified with the PolyA extensions as described in the Materials and Methods Section "Folding of DNA origami".
- **Placement of triangles**. Initial placement of the triangles was performed as described above. After placement, wafers were not dried but kept in the Placement buffer.
- **Introduction of Nanotubes.** First, the Placement buffer on a surface of the wafer was exchanged to the Hybridisation buffer with at least 8 buffer washes by pipetting 60 uL of a Hybridisation buffer onto the wafer, and pipetting 60 uL off of the wafer. DNA Origami nanotubes with polyT extensions are diluted to the concentration of 250 pM in the Hybridisation Buffer in the LowBind DNA tubes. Then, nanotubes are introduced to the buffer on the wafer by 8 buffer washes by pipetting 60 uL of the Hybridisation buffer with the nanotubes onto the wafer, and pipetting 60 uL off of the wafer. The

buffer from 8th wash is left on top of the wafer. This procedure allows to reproducibly control the concertation of the nanotubes on a surface of the wafer.

- **Incubation.** The lid of the petri dish is closed and perti dish is placed in a cell incubator, where it is incubated for 1 hour at 37°C, 98% humidity and 0% $CO_2$. After taking the wafer out of the incubator it is left for 10 minutes to cool down to room temperature.
- **Purification from excess nanotubes**. Excess origami is washed away with at least 8 buffer washes with the Hybridisation buffer followed by Tween wash. After Tween wash, the buffer on a surface of the wafer is exchanged to the Silica coating buffer.
- **Silica coating.** The wafer is placed to a well on a 12 well-plate (ThermoFisher Scentific) filled with 2.5 ml of Silica coating buffer with 4% pre-hydrolyzed silica precursors, prepared according to the protocol published in [15]. The wafer is left in this buffer for 3 days at RT.
- **Drying.** The wafer is washed in a petri dish with distilled water for 30seconds, followed by a 30 second wash in 100% Ethanol. Then the wafer is air-dried.

Notes

Importantly, we perform annealing step strictly before drying of the wafers. We noticed that after occasional dewetting or air drying from the buffer the HMDS-coated hydrophobic surface of substrates is partially deactivated. Therefore, consecutive introduction of the nanotubes in a buffer to the substrate with pre-adsorbed triangles yields high non-specific binding of the tubes to the background surrounding triangles. However, without drying background remains hydrophobic for up to 3 days at temperature up to 50$^{o}$C providing low background binding during further annealing steps.

Besides, dewetting or air drying of the wafers with annealed nanotubes (or with any other 3D DNA origami) before growth of silica shell result in collapse of the structures.

Additionally, we observed that buffer with lower $MgCl_2$ concentration of 12 mM prevents aggregation of the tubes on a surface of triangles (Supplementary Figure 13). This $MgCl_2$ concentration is still sufficient to keep triangles bound to the substrate.

**Surface annealing of DNA origami nanotubes with triangles on glass**

- **Preparation of DNA origami.** We folded and purified fresh nanotubes modified with polyT extensions and triangle modified with PolyA extensions as described in Materials and Methods Section "Folding of DNA origami".
- **Placement of triangles**. Initial placement of triangles on glass chips was performed as described in [14]. Briefly, DNA origami triangles bearing 27 Poly(A) extensions were deposited to the surface of glass chips, prepared as described in Materials and Methods Section "Nanopatterning of glass wafers with nanosphere lithography". Briefly, a 40 ml drop of origami suspension with concentration of 300 pM in Glass Placement Buffer (Supplementary Table S2) was placed on the glass chip and allowed to incubate for 1 h in a loosely capped container ringed with a moist Kimwipe to prevent excessive evaporation. After incubation, glass chips were purified from excessive DNA origami triangles by washing with 0.07% Tween-20 solution in the Glass Placement Buffer. After purification, the adsorbed DNA origami triangles could be dried in place by treating the sample with ethanol solutions, or transferred to the Hybridization buffer for the further annealing steps.
- **Introduction of Nanotubes.** First, the Placement buffer on a surface of the wafer was exchanged to Hybridisation buffer with at least 8 buffer washes by pipetting 60 uL of fresh Hybridisation buffer onto the wafer, and pipetting 60 uL off of the wafer. DNA Origami nanotubes with polyT extensions are diluted to the concentration of 1 nM in

Hybridisation Buffer in LowBind DNA tubes. Then, Nanotubes are introduced to the buffer on the wafer by 8 buffer exchanges. The buffer from $8^{th}$ wash is left on top of the wafer.

- **Incubation.** The lid of the petri dish is closed and perti dish is placed in a cell incubator, where it is incubated for 3 or 24 hours at 37°C, 98% humidity and 0% $CO_2$.
- After this step, glass chips with nanotubes are purified from excessive DNA origami, coated with the silica shell and dried as described in previous section.

**Placement of DNA origami barrels and tetrapods on Si/$SiO_2$ wafers**

- **Substrate preparation.** For the placement of DNA origami barrels, circular binding sites with diameter of 45 nm and e-beam dosage of 400 mCu/$cm^2$ arranged in square lattice with 200 nm period are defined by e-beam lithography on a Si/$SiO_2$ wafers.
  For the placement of DNA origami tetrapods, square binding sites with diameter of 35 nm with e-beam dosage of 400 mCu/$cm^2$ arranged in square lattice with 200 nm period are defined.
- **Preparation of DNA origami.** Barrels and tetrapods are folded and purified as described in the corresponding Materials and Methods Sections "Folding of DNA origami".
- **Placement**. Placement of barrels and tetrapods is performed as described above for triangles. The optimum concentration of barrels and tetrapods in the Placement buffer is 100 pM and 400 pM, respectively. After placement, the wafers are not dried but kept in the Placement buffer.
- **Silica coating.** First, the Placement buffer is exchanged to the Silica coating buffer by at least 8 buffer washes. Then wafer is incubated in Silica coating buffer with 4% pre-hydrolyzed silica precursors resulting in growth of rigid silica shell on DNA origami and dried as described in previous section.

Notes

Initially we deposited tetrapods on triangular binding sites with 50 nm size in order to obtain orientation control. However, we did not obtain control over orientation (data nor shown), mainly because of small size of the structures and small contact surface area. Therefore, we further use square binding sites to shorten the time of e-beam exposure.

**Surface annealing of DNA origami tetrapods with 24 HB on Si/$SiO_2$ wafers**

- **Substrate preparation**. For the placement of DNA origami tetrapods, square binding sites with size of 35 nm and e-beam dosage of 400 uCu/$cm^2$ arranged in honeycomb lattice with translation between next-nearest-neighbours of 170 nm are defined by e-beam lithography on a Si/$SiO_2$ wafers. Additionally, we designed a fractal arrangement of individual honeycombs forming Zelinski triangles.
- **Preparation of DNA origami.** We folded and purified fresh tetrapods and 24HBs modified with complementary extensions from both sides (as described in Materials and Methods Section "Folding of DNA origami". The tetrapods are folded without the linker extensions because they otherwise aggregate during folding.
- **Placement of tetrapods**. Initial placement of tetrapods is performed as described in the previous section. After initial placement and purification from excessive tetrapods, chips are not dried but kept in the Placement buffer.
- **Introduction of ssDNA tetrapod' linkers**
  Linker extensions (48 ssDNA staple strands extended with GGGAAGGG from 5'end) are mixed in the Placement buffer to the concentration of 40 nM. The linkers are

introduced to the Placement buffer on a surface of the $Si/SiO_2$ wafers by 8 buffer washes with 60 uL of linkers solution. The buffer from the 8th wash is left on top of the wafer and incubated for 30 mins RT. Excess linkers is washed away with at least 8 buffer washes with the Placement buffer.

- **Introduction of 24HB.** First, the Placement buffer on a surface of the wafer is exchanged to the Hybridisation buffer with at least 8 buffer washes. 24HB with 24 anchor extensions are diluted to the concentration of 1 nM in the Hybridisation Buffer in the LowBind DNA tubes. Then, 24HBs are introduced to the buffer of the wafer by 8 buffer washes. The buffer from 8th wash is left on top of the wafer.
- **Incubation.** The lid of the petri dish is closed and perti dish is placed in a cell incubator, where it is incubated for 1 hour at 37°C, 98% humidity and 0% $CO_2$. After taking the wafer out of the incubator it is left for 10 minutes to cool down to room temperature.
- After this step, wafers with networks of tetrapods and 24HBs are purified from excessive 24HBs, coated with the silica shell and dried as described in above.

**Characterization techniques**

UV-vis absorption measurements were performed with a NanoDrop ND-1000 Spectrophotometer (Thermo Scientific).

Tapping-mode dry AFM was carried out on a Dimension ICON AFM (Bruker). OTESPA silicon tips (300 kHz, Vecco Probes) were used for imaging in air. Imaged areas are 5 um x 5 um, with a resolution of 512 or 1024 pixels per line. Images are edited with the Software Gwyddeon.

TEM imaging of DNA origami lattices was carried out using a JEM-1011 transmission electron microscope (JEOL) operating at 80 or 100 kV. For sample preparation 5 ul of DNA origami diluted to 5 nM concentration were deposited on glow-discharged TEM grids (formvar/carbon-coated, 300 mesh Cu; TED Pella, Inc; prod no. 01753 - f) for 30 sec. Grids were furthermore quickly washed once with 0.1 ‰ uranyl acetate solution (5 μL) and immediately afterwards stained with 0.1 ‰ solution (5 μL) for 10 s.

SEM. The Scanning Electron Microscope used in this work is the E-line Scanning Electron Microscope (Raith). The beam settings for imaging are 10 kV acceleration and 20 um aperture.

Samples were SEM imaged after 30 s sputtering using an Edwards Sputtercoater S150B 1990. The sputter target contained 60% gold and 40% palladium. Process parameters used for sputtering were 5 mbar Ar, 1.5 kV, 11 mA. 30 s sputtering results in the deposition of layer of gold/palladium with a thickness of a few nm. SEM imaging of the samples was performed on horizontal and tilted positions by angle 70°.

**Data analysis**

We measured binding site occupancy (percentage of sites with one or more origami), number of origami at a site (0, 1, 2, or > 3), as well as origami alignment (in standing or on-site) position by hand-annotating of dry AFM (for triangular DNA origami on $Si/SiO_2$ or glass substrates) or SEM images (for DNA origami triangle-nanotubes structures, barrels and tetrapods). For all of the substrates fabricated by e-line lithography, 600 sites were analyzed with different number of independent replications *N* (experiments on the optimization of conditions of triangles-nanotubes including number of linkers, temperature and nanotubes concertation and experiments on the optimization of barrel placement are performed in single replication). Placement yields on glass substrates were counted in the middle of the glass and on 3 separated spots each 1 mm apart from the middle point. For the binding site spacing of triangles experiment, the number of sites analyzed per replicate depended on the period — 300 sites were analyzed for 170 nm and 250 nm spacing but only 144 sites were analyzed for 400 nm spacing.
